# mRNA stability in response to m^6^A placement is linked to cell identity in planarians

**DOI:** 10.64898/2026.03.27.714417

**Authors:** Constantin Höhn, Andreas Pittroff, Anne-Sophie Gribling-Burrer, Lisa Hülsmann, Redmond P. Smyth, Claus-D. Kuhn

## Abstract

N^6^-methyladenosine (m^6^A) is a prevalent internal modification of eukaryotic mRNA that influences transcript fate, including mRNA stability and cell-type-specific gene expression. However, the mechanisms underlying m^6^A-mediated regulation remain poorly understood in many systems, including the highly regenerative planarian *Schmidtea mediterranea*. To address this, we generated a high-confidence atlas of ∼72,200 m^6^A sites across the planarian transcriptome using multiplexed direct RNA sequencing. The m^6^A sites follow a DRAYW consensus motif and are highly enriched near stop codons while being largely excluded from coding sequences. This pattern aligns with an exon length-dependent variant of the exon junction complex-mediated (EJC) exclusion model, wherein the EJC restricts m^6^A deposition near splice sites. Knockdown of the m^6^A writer complex induced pronounced, cell-type-specific changes in transcript stability. Destabilized transcripts were enriched for intestinal markers, whereas stabilized transcripts were associated with neoblasts, the adult stem cells of planarians. Transcriptional shut-off experiments further confirmed that m^6^A has opposing effects on mRNA decay depending on cellular context: it stabilizes transcripts in differentiated cells, while it promotes the degradation of mRNAs associated with neoblasts. Collectively, these results support a model in which cell-type-specific regulation of mRNA stability by m^6^A plays a crucial role in shaping cell identity in planarians.

**GRAPHICAL ABSTRACT:** 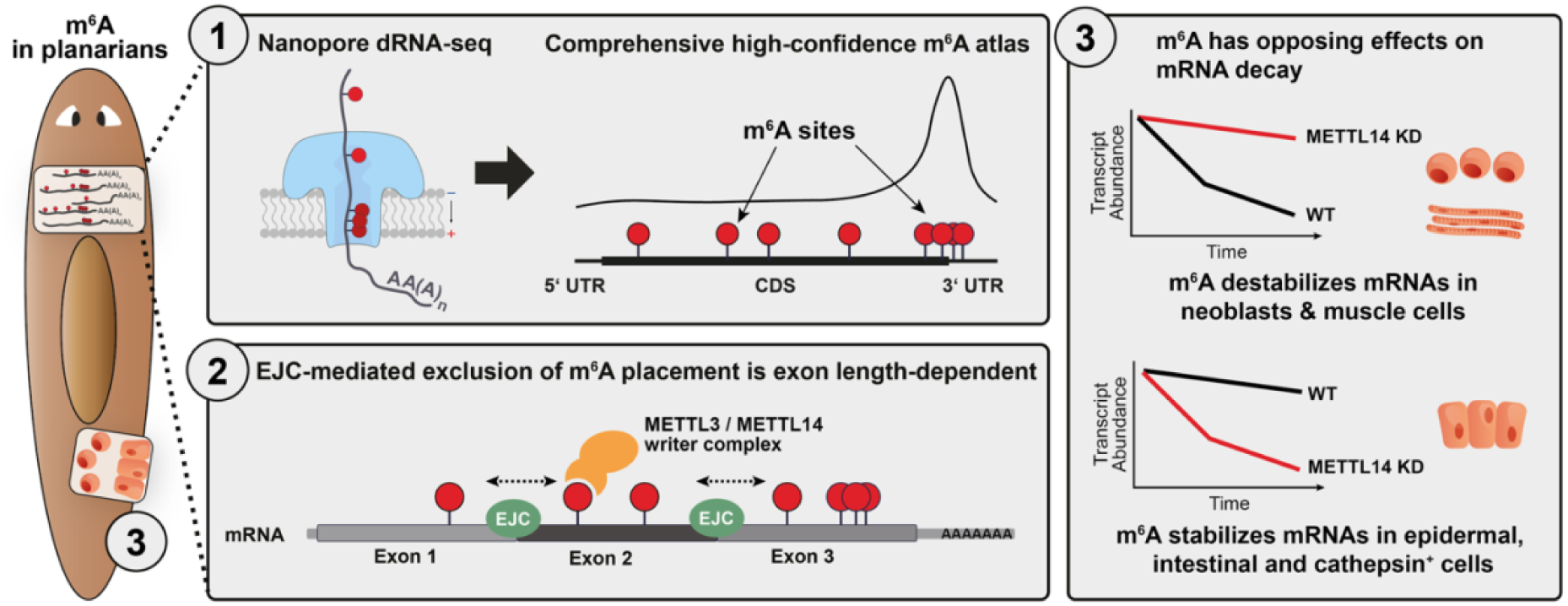

## INTRODUCTION

The interest in modified nucleobases in mRNAs has seen a tempestuous renaissance in the past decade. In contrast to the omnipresence and functional importance of modified nucleobases for transfer RNAs (tRNAs) and ribosomal RNAs (rRNAs), modifications on mRNAs were only recently found to possess essential and diverse functions in almost all eukaryotic organisms (1, 2). Out of >150 possible types of modifications detected so far, N^6^-methyladenosine (m^6^A) is by far the most prevalent, with 0.15 – 0.6% of all adenosines being methylated in human cells (3). m^6^A modifications require a sophisticated machinery for their installation and for mediating downstream functions. First, a large methyltransferase complex (MTC), which forms in the nucleus around the catalytic core of the METTL3-METTL14 heterodimer, installs m^6^A on mRNAs (4). Second, a diverse set of both nuclear and cytoplasmic m^6^A reader proteins, predominantly from the YTH domain family of proteins, recognizes m^6^A-modified mRNAs and orchestrates downstream functions (5). Lastly, m^6^A demethylases, such as ALKBH5 and FTO in humans, reverse m^6^A placement by actively removing it from mRNAs (Shen et al., 2022). The molecular functions of m^6^A do not appear to be limited to a single mode of action. In contrast, they are diverse and range from affecting mRNA half-life (7, 8), at times in conjunction with translation (9), to altering mRNA splicing (10), promoter-proximal pause release (11) and translation efficiency (12).

The mode of action of m^6^A-modified adenosines is primarily determined by three factors: (i) the intracellular localization and identity of the m^6^A reader that mediates their effects; (ii) the precise position of the modification within individual mRNAs (7, 9); and (iii) m^6^A-induced remodeling of RNA structure, which influences downstream regulatory processes and reshapes the RNA interactome (13, 14). In *D. melanogaster*, for example, m^6^A is enriched at the 5’ ends of transcripts. Consequently, the MTC, in conjunction with the reader YTHDC1, was found to regulate the release of paused RNA polymerase II (Pol II), thereby significantly affecting mRNA synthesis rate (11). In contrast, human transcripts exhibit low - but nonetheless significant - levels of m^6^A across their entire open reading frame, with a characteristic enrichment of m^6^A around the stop codon (15). In humans, the MTC installs m^6^A specifically on adenosines located within so-called DRACH motifs (where D = A/G/U, R = A/G, A = the methylated adenosine, C = cytosine, and H = A/C/U) (16). However, despite the abundance of DRACH motifs in the human transcriptome, only a fraction of these is methylated. Selecting the adenosines that are to be methylated in coding sequences was found to depend on the exon junction complex (EJC), which serves as a physical obstacle to m^6^A placement close to exon-intron junctions in human cells. All other adenosines that are at the center of suitable DRACH motifs are equally likely methylated by the MTC (7). However, it is unknown whether this model of m^6^A placement holds true for all higher organisms, in which mRNAs were found to carry m^6^A modifications.

To tackle this question, we explored the influence of m^6^A on mRNA surveillance in the highly regenerative flatworm *Schmidtea mediterranea*, a popular model organism for studying whole-body regeneration fueled by pluripotent stem cells (17). In *S. mediterranea*, roughly 24% of all transcripts carry at least one m^6^A modification, with 0.5% of adenosines being methylated (18, 19). This modification plays a critical role in planarian regeneration, tissue homeostasis, stem cell differentiation - particularly in the intestinal lineage - and cell cycle regulation (19) (18). Recent single-nucleotide resolution mapping using GLORI (15, 20) identified 19,328 high-confidence m^6^A sites (>10% methylation) in the planarian transcriptome (21). While m^6^A marks are installed on adenosines within a degenerate sequence motif akin to the human DRACH motif, an expanded family of YTDHF m^6^A reader proteins forms a robust and redundant network critical for proper lineage specification in planarians (Yesharim et al., 2026).

Yet, despite progress in understanding the impact of m^6^A on stem cell differentiation and lineage commitment, its direct influence on mRNA stability and surveillance remains unexplored. This study addresses this knowledge gap, providing novel insights into the role of m^6^A in mRNA regulation in planarians. First, we examined the effects of MTC depletion on transcript levels and evaluated these changes considering the exact position and number of m^6^A sites per transcript. This analysis revealed cell type-specific gene expression changes, where loss of m^6^A was mainly linked to transcript downregulation, largely independent of where m^6^A sites were located. Second, we studied the impact of m^6^A on transcript stability after blocking planarian transcription by Actinomycin D, which revealed that m^6^A exerts opposing effects on mRNA stability depending on cellular context. Whereas transcripts in cathepsin+, epidermal and intestine cells were stabilized, we found mRNA decay to be promoted in neoblasts and muscle cells. Importantly, we based our results on precisely mapped m^6^A sites using Nanopore direct RNA sequencing (dRNA-seq). This approach allows for the determination of m^6^A sites with single-nucleotide precision. Moreover, as dRNA-seq does not rely on the chemical conversion of non-modified adenosines and additional reverse transcription steps, as does GLORI (21), the data presented in this study constitute both the most complete and the most reliable dataset of planarian m^6^A sites available to date. Using this comprehensive dataset in combination with EJC depletion in planarians, we demonstrate that the EJC restricts MTC access to m^6^A sites proximal to exon-exon junctions, as previously shown for human cells (7). However, in planarians we found the exclusion zone for m^6^A placement to depend on exon length and to be shifted slightly downstream. Our analysis further revealed a pronounced enrichment of m^6^A sites near stop codons, while only few sites are present throughout the coding regions of planarian transcripts, in contrast to comparably higher levels of coding sequence (CDS)-linked sites in humans. Collectively, our results reveal that m^6^A is an essential, direct actor in determining the life cycle of marked transcripts in planarians. Together with the recent study on the effect of m^6^A reader protein on proper lineage specification (21), this study paves the way to decipher the interconnection between m^6^A-marked transcripts, m^6^A reader proteins and proper stem cell differentiation.

## MATERIAL AND METHODS

### Animal Husbandry

The asexual *Schmidtea mediterranea* (clonal line CIW4; Sánchez Alvarado et al., 2002) was maintained in darkness at 20 °C in 1x Montjuic medium (1.6 mM NaCl, 1 mM CaCl_2_, 1 mM MgSO_4_, 0.1 mM MgCl_2_, 0.1 mM KCl, and 1.2 mM NaHCO3). Animals were fed once a week with homogenized calf liver. For experimental procedures, planarians were kept in small boxes filled with 1x Montjuic medium supplemented with 50 µg/mL gentamycin sulfate and maintained under the same conditions (20 °C, darkness). Prior to experiments, worms were starved for a minimum of 7 days.

### RNAi Knockdowns

Inserts carrying the target sequences for *mettl3*, *mettl14*, *eIF4A3*, *y14* knockdowns were amplified from planarian cDNA and cloned into the pPR-T4P vector via Gibson assembly (Supplementary Table S1) (Gibson et al., 2009). Double-stranded RNA (dsRNA) for RNAi was generated as previously described (24). Briefly, the templates for *in vitro* transcription (IVT) were amplified by Taq-based PCR from plasmids containing the cloned target sequence. PCR products were purified with the QIAquick PCR purification Kit (Qiagen) according to the manufacturer’s protocol. dsRNA for knockdown experiments was generated by T7 RNA polymerase-mediated *in vitro* run-off transcription (25, 26). The two T7 promoters flanking the fragment of interest enable the production of dsRNA. IVT reactions were prepared in 150 µL and incubated at 37 °C at 600 rpm for at least 4 h. Following transcription, 17.1 µL DNase I incubation buffer and 1.5 µL DNase I were added to each reaction, and the mixture was incubated for 15 min at 37 °C and 800 rpm. Reactions were terminated by adding 19.2 µL 500 mM EDTA (pH 8.0) to a final concentration of 50 mM, followed by vortexing. Product dsRNA was purified by precipitation with three volumes of 100% ethanol and a final concentration of 0.3 M sodium acetate. After resuspension in RNAse-free ddH_2_O, dsRNA was formed by heating to 90°C for 3 min, followed by gradual cooling to 25 °C at a rate of 0.1°C/s. Residual nucleotides were removed using Microspin G50 columns (GE Healthcare) following the manufacturer’s instructions.

For RNAi experiments, worms were starved for at least 7 days prior to treatment and fed twice a week with calf liver mixed with a total of 3 µg dsRNA per animal. When multiple dsRNAs were fed, the amount of each individual dsRNA was adjusted accordingly. Control dsRNA (*unc22*) was included in knockdown experiments to monitor non-specific effects of the RNAi treatment.

### Actinomycin D assay

Knockdowns targeting *mettl14* and *unc22*, along with wildtype controls, were established and maintained until 22 dpff. For each condition, 40 worms were fed twice a week with either dsRNA-mixed calf liver (experimental groups) or calf liver alone (wild-type controls). At 22 dpff, worms were transferred to Petri dishes containing 1x Montjiuc medium supplemented with actinomycin D (final concentration: 200 µg/mL). At 0, 6, and 12 h post-actinomycin D treatment, three worms per biological replicate were collected (four replicates per time point). Worms were transferred to 1.5 mL tubes containing 100 µL of TRIzol, snap-frozen in liquid nitrogen and stored at -80 °C until RNA extraction and library preparation. Time points were selected based on preliminary experiments indicating detectable transcript decay after 6 hours (not shown).

### RNA extraction for qPCR samples

To extract RNA for qPCR, one or three planarians were homogenized in 100 µL TRIzol using a douncer. The volume was adjusted to 500 µL with additional TRIzol. 100 µL chloroform was added per 500 µL of TRIzol, and the mixture was mixed by inverting it. After incubating for 3 min at room temperature, samples were centrifuged for 15 min at 4 °C with 12000x g. The upper aqueous phase, containing RNA, was transferred to a new tube. Subsequently, 1:1 volume of 100% ethanol was added to the aqueous phase, and the mixture was applied to a Zymo-Spin^TM^ column. The column was centrifuged at 16000x g for 30 s and the flow-through was discarded. The column was washed with 400 µL of RNA Wash Buffer and centrifuged again for 30 s at 16000x g. For on-column DNase treatment, 0.5 μL of DNase I (10 U/μL) and 39.5 μL of 1x DNA Digestion Buffer (10x DNA Digestion buffer: 100 mM Tris-HCl pH 9.5, 25 mM MgCl_2_, 1 mM CaCl_2_; 1x DNA Digestion buffer: 100 µL 10x DNA Digestion buffer, 700 µL 100% ethanol, 200 µL DEPC-treated ddH_2_O) were added to the column. The mixture was incubated at room temperature for 15 min. After incubation, the column was centrifuged at 16000x g for 90 s and the flow through was discarded. The column was then washed with 400 μL of RNA Preparation Buffer, centrifuged at 16000x g for 30 s, and the flow-through discarded. Next, 700 μL of RNA Wash Buffer was added, the column was centrifuged at 16000x g for 2 min, and the flow-through was discarded. This wash step was repeated with 400 μL of RNA Wash Buffer and centrifuged for 1 min. To remove residual ethanol, the column was centrifuged again at 16000x g for 1 min. RNA was eluted in 10 - 15 μL of DEPC-treated water by incubating the column for 1 min at room temperature, followed by centrifugation at 16000x g for 90 s.

### Large-scale RNA extraction for Nanopore dRNA-Seq

For large-scale RNA extraction, 35 fully grown worms (MTC RNAi experiment; animals had been fed twice weekly for 4 weeks) or 20 fully grown worms (EJC RNAi experiment; animals had been fed once (6 dpff) or twice (8 dpff)) were picked and transferred into a medium-sized homogenizer tube. 10 mL of TRIzol was added, and the mixture was homogenized using a douncer. An additional 10 mL of TRIzol was added, and the mixture was homogenized again. The lysate was divided equally between two 20 mL centrifuge tubes. To maximize RNA yield, another 10 mL of TRIzol was added to the homogenizer tube, and the residual lysate was recovered by pipetting up and down. This additional volume was split between the two centrifuge tubes. The mixture was incubated for 5 min at room temperature. Chloroform (3 mL) was added to each tube, and the contents were mixed by pipetting up and down. After incubation for 3 min at room temperature, the samples were centrifuged for 15 min at 4 °C with 12000x g. The upper aqueous phase was transferred to a 50 mL falcon tube. Extracted RNA was purified using the Zymo RNA Clean & Concentrator Kit 25. The volume of extracted RNA solution was measured using a pipette controller. 1 volume of 100% ethanol was added to 1 volume of extracted RNA. The mixture was divided across 16 Zymo-Spin^TM^ columns (each in a 2 mL collection tube). Due to the column’s maximum load volume of 800 µL, each column was loaded three times. On-column DNAse treatment and purification were performed as described above, but the DNAse reaction mix was scaled up twofold. Each column received 80 µL of the DNase mixture. RNA was eluted in 32 µL DEPC-treated water.

### RNA extraction of actinomycin D assay samples

To extract RNA from actinomycin D-treated samples, three worms per replicate were homogenized as described above. TRIzol (900 µL) was added to each sample, and the mixture was incubated at room temperature for 5 min. Chloroform (200 µL) was added, and the mixture was incubated again. Samples were centrifuged for 15 min at 4 °C and 12000x g. The aqueous phase was transferred to a new tube, and an equal volume of 100% isopropanol was added. After incubation for 10 min at room temperature, the precipitate was pelleted by centrifugation for 10 min at 4 °C and 12000x g for 10 min. The RNA pellet was subsequently washed with 1 mL of 70% ethanol and resuspended in 43 µL DEPC-treated water. DNAse treatment (43 µL RNA solution, 5 µL 10x DNAse buffer, 2 µL DNAse I (10 U/µL)) was performed for 25 min at 37 °C. To size-select for RNAs > 200 nt, 100 µL 2x RNA Binding buffer (Zymo-Research) and 50 µL 100% ethanol were added to the DNase-treated RNA. The mixture was applied to a Zymo-Spin^TM^ IICR column, and the cleanup was performed as described above. RNA was eluted in 30 µL of DEPC-treated water.

### cDNA synthesis and real-time qPCR

cDNA was synthesized from 1-5 µg total RNA using SuperScript IV Reverse Transcriptase, according to the manufacturer’s instructions. The efficiency of knockdown experiments and the reduction of target transcripts were assessed by Real-time quantitative PCR (RT-qPCR).

Amplification was performed using a CFX Connect^TM^ Real-Time System, and data were analyzed with Bio-Rad CFX Manager^TM^ software (v2.1). Primers used for qPCR are listed in Supplementary Table S1. RT-qPCR reactions were performed in technical triplicates, and relative expression was calculated using the Pfaffl method (27).

### mRNA isolation

Poly(A) mRNA was enriched using the NEBNext^®^ Poly(A) mRNA Isolation Module for m^6^A specific mass spectrometry. mRNA isolation was performed according to the manufacturer ’s instructions. The buffers (2x Binding buffer: 0.5 M LiCl, 0.1 M Tris-HCl (pH 8.0), 0.01 M EDTA, 1% (w/v) Lithium dodecylsulfate, 5 mM DTT and Washing buffer: 0.15 M LiCl, 0.01 M Tris-HCl (pH 8.0), 1 mM EDTA, 0.1% (w/v) SDS) were prepared as described (28). For Nanopore sequencing, the NEBNext^®^ High Input Poly(A) mRNA Isolation Module was used according to manufacturer’s instructions. In deviation from the standard protocol, elutions were performed at 70 °C to preserve the integrity of longer RNA transcripts.

### rRNA depletion

rRNA depletion was performed using the siTOOLs Biotech riboPOOL system, which is based on prior work from our lab (29). To 14 µL of RNA, 1 µL riboPOOL and 5 µL of Hybridization Buffer (siTOOLs) were added. The mixture was incubated at 68 °C for 10 minutes in a heating block, followed by 30 minutes at 37 °C. Meanwhile, the streptavidin-coated magnetic beads were prepared. For each sample, 90 µL of bead suspension were transferred to a reaction tube and placed on a magnetic rack until the solution cleared. The supernatant was removed, and the beads were resuspended in 120 µL Depletion Buffer per sample. The bead suspension was split into two tubes (tube 1 and tube 2). After placing the tubes on the magnetic rack and removing the supernatants, the beads in tube 1 were resuspended in 220 µL Depletion buffer, and the beads of tube 2 were resuspended in 120 µL depletion buffer per sample. To capture rRNA-DNA hybrids, 220 µL of beads from tube 1 were added to the hybridized RNA sample and incubated for 15 min at 37 °C. Tubes were tapped every 3 min to keep the beads in suspension. The tube was then placed on the magnetic rack, and the supernatant was transferred to a new tube. An additional 120 µL of beads from tube 2 were added to the supernatant, followed by another incubation for 15 min at 37 °C. After placing the tube on the magnetic rack, the supernatant was mixed with 360 µL 2x RNA Binding buffer (Zymo-Research) and 360 µL 100% ethanol. The entire binding mixture was loaded onto a Zymo-IC column and RNA purification was performed as described previously (RNA extraction for qPCR samples). In deviation from the standard protocol, the on-column DNAse treatment mixture was performed using 1 µL DNase I (recombinant, RNase-free; Roche, 10 U/µL), 3 µL 10× DNase I Buffer (Roche) and 26 µL RNA Wash Buffer (Zymo-Research).

### Mass spectrometric analysis

To quantify alterations in m^6^A levels on mRNAs following knockdown experiments, poly(A) selected RNA was analyzed by Cathrine Broberg Vågbø, PhD. at the Proteomics and Modomics Experimental Core Facility (PROMEC) of the Norwegian University of Science and Technology (NTNU). Statistical analysis of the resulting data was performed using the Tukey-HSD test with p-values adjusted for multiple comparisons.

### RNA Library preparation and NovaSeq sequencing

Library preparation and sequencing of the actinomycin D assay was performed from rRNA depleted RNA by Genewiz Germany GmbH. RNA samples were quantified using Qubit 4.0 Fluorometer (Life Technologies, Carlsbad, CA, USA) and RNA integrity was checked with RNA Kit on Agilent 5300 Fragment Analyzer (Agilent Technologies, Palo Alto, CA, USA). RNA sequencing libraries were prepared using the NEBNext Ultra II Directional RNA Library Prep Kit for Illumina (NEB, Ipswich, MA, USA) according to the manufacturer’s general workflow. Briefly, ribo-depleted RNA was taken directly into the fragmentation and first-strand synthesis module. First-strand cDNA was generated using random priming, and second-strand synthesis was performed in the presence of dUTP to enable strand specificity through downstream degradation of the second strand during amplification. Following double-stranded cDNA generation, fragments underwent 3′ end-repair and adenylation. Adapters containing unique molecular identifiers (UMIs) were ligated to the A-tailed cDNA fragments. After adapter ligation, libraries were amplified using limited-cycle PCR to enrich adapter-ligated molecules and selectively degrade the dUTP-containing second strand, preserving library directionality. Completed libraries were assessed for fragment size distribution using an NGS fragment analysis system (e.g., Agilent 5300 Fragment Analyzer; Agilent Technologies, Palo Alto, CA, USA) and quantified using a fluorescence-based assay (e.g., Qubit 4.0 Fluorometer; Invitrogen, Carlsbad, CA, USA). The sequencing libraries were multiplexed and loaded on the flow cell on the Illumina NovaSeq X Plus instrument according to manufacturer’s instructions. The samples were sequenced using a 2x150 Pair-End (PE) configuration v1.5. Image analysis and base calling were conducted by the NovaSeq Control Software on the NovaSeq instrument. Raw sequence data (.bcl files) generated from Illumina NovaSeq was converted into fastq files and de-multiplexed using Illumina bcl2fastq program version 2.20. One mismatch was allowed for index sequence identification. After investigating the quality of the raw data, sequence reads were trimmed to remove possible adapter sequences.

### Nanopore direct RNA sequencing and base calling

Nanopore direct RNA sequencing was performed following the WarpDemuX adapter-barcoding and demultiplexing strategy as described (30). Briefly, custom RTA adapters (10 µM forward, 10 µM reverse) were annealed in 30 mM HEPES-KOH (pH 7.5), 100 mM K-Acetate by incubation for 1 min at 95 °C followed by a slow cooling to 25 °C at a rate of 0.5 °C/min. The annealed RTA were then diluted to 0.7 µM and stored at -20 °C until use. For each sample, 100 ng of polyA-selected RNA were ligated to 1 µL of 0.7 µM annealed RTA in the presence of 2 µl 5x NEBNext Quick Ligation Buffer (NEB), 0.5 µl T4 DNA Ligase high concentration (NEB) and 0.2 µl RNAsin (Promega) in a final volume of 10 µL by incubation at 22°C for 15 min. To stop the ligation, 2 µl of 0.5 M EDTA was added and samples were pooled and purified with 0.5 volumes of SPRI beads (Mag-Bind TotalPure NGS, Omega Bio-tek), washed twice with 200 µl fresh 80 % EtOH, and eluted in 10 µl RNase-free H2O. Reverse transcription was performed by adding 8 µl of 5X Induro buffer, 2 µl of 10 mM dNTP, 2 µl of 10 uM random hexamer oligonucleotides, 2 µl Induro RT (NEB), 0.5 µl RNasin in a final volume of 40 µl and incubation 2 min at 22°C (annealing), 15 min at 60°C (reverse transcription) and 10 min at 70°C (heat-inactivation). The reaction was purified with 1.4 volumes of SPRI beads, washed twice with 200 µl fresh 80 % EtOH, and eluted in 10 µl RNase-free H_2_O. The sequencing adapter ligation was realized using the SQK-RNA004 library kit (ONT) according to the manufacturer’s instruction. Briefly, the 10 µl purified library was mixed with 6 µl 5x NEBNext Quick Ligation Buffer, 5 µl RLA sequencing adapter (ONT SQK-RNA004) and 3 µl T4 DNA Ligase high concentration in a final volume of 30 µl and incubated 15 min at 22°C. The reaction was purified with 0.5 volumes of SPRI beads, washed twice with 100 µl RNA wash buffer (WSB, ONT SQK-RNA004). The library was then eluted in 32 µl RNA elution buffer (REB, ONT SQK-RNA004). The library was loaded onto a Promethion (FLO-PRO004RA) flow cell for sequencing. Data acquisition was performed using MinKNOW version 24.11. Basecalling was performed with dorado v0.8.3 using the model parameters sup v5.0.0 m^6^A with poly(A) length estimation. Demultiplexing was performed using the demux command of WarpDemuX v0.4.5.

### Processing, mapping and m^6^A calling of Nanopore dRNA-Seq data

All genome and transcriptome reference files (version schMedS3) used in this study were retrieved from (31). Sequences from demultiplexed Nanopore dRNA-Seq libraries were extracted using samtools fastq (32). Prior to mapping, reads were aligned to a combined reference database comprising *S. mediterranea* rRNA sequences (29) and tRNA sequences (GtRNAdb; http://gtrnadb.ucsc.edu). Unmapped reads were filtered using the samtools flag -f 0x4 and the resulting rRNA- and tRNA-depleted reads were then mapped to the planarian genome. Both mapping steps were performed using minimap2 (-ax map-ont -uf -k 14; -ax splice -uf -k 14 respectively; (33, 34). Mapped reads were filtered using the samtools flag -F 0x4 to retain primary and secondary alignments for downstream analysis. m^6^A modification probabilities and levels were assessed using modkit’ sample-probs function (Modkit, 2023/2026). To construct bedMethyl tables, genome-mapped reads were processed with modkit’s call-mods and pileup utilities (modification threshold = a:0.99; filter-threshold 0.9; Modkit, 2023/2026). The resulting bedMethyl tables were filtered to retain m^6^A calls covered by ≥10 reads, and ≥15% methylation rate. Calls were compared across replicates and only those fulfilling these criteria in at least two of three replicates were retained for further analysis (Supplementary Table S2).

### Motif assessment and m^6^A distribution

Metaplots of m^6^A distribution were generated using the R package metaplotR (35). GenePred files were derived from the planarian trancriptome (version schMedS3; (31)) using UCSC’s gtftoGenePred utility (https://github.com/BioQueue/ucsc-tools). To investigate the sequence context of m^6^A installation, previously identified m^6^A sites were expanded by two nucleotides in each direction using bedtools slop (-b 2). The resulting 5-mer sequences were extracted with bedtools getfasta (36). Nucleotide distributions within these 5-mer sequences were assessed, and nucleotides with a contribution exceeding 20% at any individual position were considered as part of the m^6^A installation consensus. Sequence motifs were visualized in R using the Biostrings (37), seqLogo (38), and DiffLogo (39) packages. The *S. mediterranea* genome was scanned for all possible 5-mer combinations of nucleotides within the consensus motif using patman (40). Genic motif occurrences were intersected with bedMethyl tables using bedtools intersect (36) to assess the distribution of methylation rates for each individual 5-mer.

### Differential Gene Expression Analysis

Nanopore dRNA-Seq reads were mapped to the planarian transcriptome (31) using minimap2 (-ax map-ont -k 14 -N 10; (33, 34)) and quantified with salmon (salmon quant --ont --noLengthCorrection -l A; (41)). Resulting transcript level abundance estimates were summarized to gene level using tximport (42). Differential gene expression analysis was performed using DESeq2 (43) with the design formula ∼condition. All possible comparisons were calculated: *metll14* (RNAi) vs wildtype, *unc22* (RNAi) vs wildtype, *mettl14* (RNAi) vs *unc22* (RNAi). Genes with a log2 fold change exceeding ±0.5 and an adjusted p-value < 0.05 were considered significantly deregulated. Log_2_ fold changes were shrunk using DESeq2’s lfcShrink function (type = “normal”). Volcano plots were generated using the EnhancedVolcano package (44). Additionally, we calculated the log_2_ fold change of gene-wise m^6^A site abundance to reflect differential gene expression with respect to respective m^6^A changes:

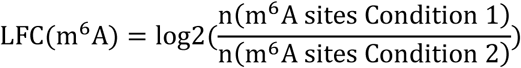

### Cell identity mapping for genes of interest

Single cell expression data were downloaded from the planarian single-cell atlas (45). To generate a reference dataset of cell-type specific markers, we selected transcripts associated with only one distinct cell type. Transcripts with ambiguous or multi-tissue associations were excluded from the reference dataset.

Since the planarian single-cell atlas is based on an older version of the planarian transcriptome version, we converted dd_v6 transcript IDs to schMedS3 transcript IDs to enable comparison with our dataset. To achieve this, dd_v6 transcripts were mapped to the schMedS3 transcriptome using minimap2 (-x splice -k 14 -N 10; (33, 34)). The resulting PAF file was filtered for sense hits with ≥ 80% match rate. Secondary alignments were retained only for isoforms of the primary alignment. This process allowed us to assign dd_v6 IDs to ∼50.78% of schMedS3 transcripts. Where possible, schMedS3 transcripts IDs were converted to dd_v6 IDs for genes of interest and compared to the reference database of cell-type-specific markers. Overrepresentation and depletion of cell types was determined using an odds-ratio, with trends categorized as biologically meaningful if the relative change in representation exceeded 0.3.

### Methylation frequency at exon-intron junctions

Exons in the *S. mediterranea* transcriptome were classified as single, first, internal, or last exons based on annotated transcripts, accounting for multiple definitions of the same exon across different isoforms. m^6^A sites and motif occurrences were overlapped with these exon annotations using bedtools intersect (36). m^6^A-harboring exons were then stratified into four length classes based on exon length quartiles. The distance of m^6^A sites and motifs to the respective exon boundaries was calculated in bins of 10 nt. To model methylation frequencies as a function of distance to internal exon starts and ends, as well as last exon starts, generalized additive models (GAMs) with a binomial distribution and logit link function were fitted separately for each exon length class using the gam() function from the R package mgcv (46). The models had the following form (in R formula notation): cbind(Sites, Motifs-Sites)∼s(Distance, k=6)

Here*, Sites* represents the number of observed m^6^A sites per 10-nt bin, *Motifs* represents the total number of detected motif occurrences in the same bin (as described in section Motif assessment and m^6^A distribution), and *Distance* is the distance from the exon junction of the corresponding bin (each consisting of 10 nucleotides). The response term *cbind(Sites, Motifs − Sites)* encodes the number of successes (methylated motifs) and failures (unmethylated motifs) for the binomial model. The smooth term *s(Distance, k = 6)* allows a flexible, non-linear relationship between methylation probability and distance from the exon boundary while limiting the spline complexity to six basis functions. For calculation of continuous generalized additive models, we merged the nucleotide bins up- and downstream of the exon splice junction, creating a total of 100 (-50 to +50) bins of 10 nucleotides each. Models for human data were modelled in the same way.

### Bayesian mixture model for the effect of m6A position on differential gene expression upon m^6^A loss

We fitted a Bayesian hierarchical mixture-of-experts model to transcript-level log_2_ fold change estimates of gene expression upon m6A loss, with three components representing negative, null, and positive regulatory effects. Observations were modeled using Student-t likelihoods with component-specific means and a scale parameter given by sqrt(se^2 + sigma^2), combining known measurement error with an estimated residual variance term. The student-t degrees of freedom parameter was estimated from the data, allowing tail heaviness to adapt flexibly. The negative and positive component means were parameterized as signed softplus-transformed offsets beyond a minimum effect size of 0.5, applied to linear predictors in m^6^A position, ensuring directional consistency while allowing effect magnitude to vary smoothly with m^6^A position along transcripts. A hierarchical prior linked the positive and negative component magnitude parameters through a shared location-scale structure, providing partial pooling while preserving biologically interpretable sign constraints. The null component mean was strongly regularized around zero. Mixture probabilities were modeled through a multinomial logistic gating function with the null component as the reference category, allowing regulatory state probabilities to vary with m6A position. The slopes governing position dependence of the gating probabilities were strongly regularized to reduce confounding between changes in mixture membership and changes in component-specific effect magnitudes. We included all transcripts that lost at least 80% of their m6A sites, defined as those with a methylation rate of at least 15% before and 0% after METTL14 knockdown (n = 3,966). For each transcript, m6A position was defined as the average position of all lost m6A sites within that transcript.

### Poly(A) tail analysis from Nanopore dRNA-Seq data

Poly(A) tail status and length was assessed by parsing the pt:i: flag from Nanopore dRNA-Seq reads. The pt:i: flag reports the number of nucleotides in the poly(A) tail, as estimated by Nanopore’s basecalling algorithm. To analyze poly(A) tailing in distinct transcript or gene clusters, reads overlapping these features were identified using bedtools intersect (36). Read-IDs from these overlaps were extracted and used to select the corresponding Nanopore dRNA-Seq reads for further analysis.

### Processing and Mapping of Actinomycin D assay sequencing data

Adapters were trimmed from paired-end reads using trimmomatic in paired-end mode to retain synchronized reads (ILLUMINACLIP:2:30:10, SLIDINGWINDOW:5:20; (47)). rRNA and tRNA reads were filtered using SortMeRNA and bowtie2, respectively ((29, 48–50); http://gtrnadb.ucsc.edu). Filtered reads were mapped to the planarian transcriptome (version schMedS3; (31)) using bowtie2 (--no-unal, --very-sensitive, --no-mixed; (50)). PCR duplicates introduced during library preparation were collapsed using umicollapse (51). Transcriptome-mapped and UMI-collapsed reads were quantified using salmon (41).

### Analysis of transcript decay rates in Actinomycin D assay

Salmon quantifications of transcriptome-mapped sequencing data were imported and summarized to gene-level counts using tximport (42). Counts were normalized to untransformed counts per million (CPM) using edgeR (52). Since *mettl14* dsRNA fragments were detected in the sequencing data, *mettl14* gene counts were removed prior to normalization. Normalized gene counts were filtered to retain only m^6^A-harboring genes. Linear models of gene abundance over the time course of the Actinomycin D assay were computed for each condition separately (gene expression ∼ time). Log_2_ fold changes between the 0-hour and 6-hour or 12-hour time points were calculated for each gene. P-values from the linear models were adjusted for multiple testing using the Benjamini-Höchberg procedure. To reduce noise, genes displaying a positive slope in gene abundance (p-adj ≤ 0.05) and a log2 fold change ≥ 0.2 over the time course were excluded from further analysis. CPM values were centered to the 0-hour time point for each individual gene using the following formula:

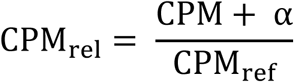

where CPM is the counts per million at the respective timepoint, α is a pseudocount (0.0001) to avoid division by 0, CPM_ref_ = CPM_0h_ + α, resulting in relative CPM values. Using maSigPro (53), we defined a quadratic regression model (degree = 2) for the time course experiment to capture non-linear decay dynamics over time. We identified significant genes using the p.vector function (Q = 0.01, MT.adjust = “BH”, min.obs = 20). After removing influential genes as defined by the T.fit function, stepwise regression was performed using the two-ways forward method at a significance level of 0.05.

The optimal number of clusters for the differential abundance profile time series analysis was estimated using within-cluster sum of squares (WCSS), silhouette analysis, and principal component analysis (PCA). Time series analysis was conducted and genes with significantly differential gene abundance profiles between *unc22* (RNAi) and *mettl14* (RNAi) conditions were selected using maSigPro’s see.genes function (show.fit = T, cluster.method = “hclust”, cluster.data = 1, k = 2; (53))

## RESULTS

### A high-confidence planarian m^6^A atlas derived from multiplexed Nanopore direct RNA sequencing

When analyzing the most recent version of the planarian transcriptome (31) for components of the m^6^A machinery, we identified core components of the MTC, as well as numerous m^6^A reader proteins of the YTH and HNRNP families via BLAST, in line with previous reports (19, 21) (Supplementary Table S3). Notably, and as is the case in *D. melanogaster* (54), we found no evidence of m^6^A demethylases (“erasers”) in the planarian transcriptome. While this suggests that m^6^A marks may not be actively removed in planarians, we cannot exclude the possibility that uncharacterized or divergent readers exist. Previous studies have demonstrated that the YTHDF reader family is expanded in planarians (21), which, together with the *bona fide* MTC (with METTL3 and METTL14 at its core), suggests that m^6^A is installed on planarian mRNAs and subsequently read by a diverse set of m^6^A readers to exert numerous biological functions (18, 19, 21).

To establish the basis for investigating the direct effect of m^6^A on planarian transcripts, we applied dRNA-seq to polyA-selected mRNA isolated from whole animals. We did so to detect and characterize all m^6^A sites that are robustly methylated in the transcriptome of *S. mediterranea* with single nucleotide precision. To minimize batch effects and other sources of error, we utilized a novel protocol that allows multiplexing of up to six Nanopore dRNA-seq libraries into one sequencing run (Fig. 1A) (30). Using this method, we obtained three dRNA-seq datasets averaging ∼5.9M reads with a median read length of 464 nucleotides (nt) and N50 value of 687 nt (Supplementary Table S4). Transcriptome coverage was extensive, as we were able to detect corresponding direct RNA reads for 18,095 (84.6%) of 21,401 genes in the planarian genome in at least one replicate, and for 14,809 (69.2%) in all three replicates. Oxford Nanopore’s Dorado software was used to directly detect N^6^-methyladenosines during base calling. This procedure resulted in a robust base modification profile, which allowed us to use a stringent cutoff of 99% base modification probability for the calculation of bed-methyl tables with modkit (55) (Fig. 1B). Next, we analyzed the methylation rate of individual m^6^A sites, which revealed that most sites exhibit methylation levels below 15%, a finding that becomes even more pronounced at base modification probability thresholds of 60% or 80% (Fig. 1C). Consequently, we excluded these lowly methylated sites from all subsequent analysis. Moreover, because low methylation rates could severely impede connecting observed transcript changes to m^6^A methylation, we decided to only consider sites with methylation rates ≥15% and required every m^6^A site to meet this threshold in at least two out of three replicates for further analysis (red dashed bar in Fig. 1C). This threshold mimics the cutoff applied to the gold standard human GLORI data (15) and is more stringent than presented for planarians before (21). At a methylation threshold of 15%, our analysis identified 72,237 high confidence m^6^A sites distributed over 28,221 transcripts and 12,583 genes respectively, rendering this dataset the most comprehensive m^6^A resource for planarians to date (Supplementary Table S2 lists genomic coordinates of all sites meeting this threshold with their respective methylation levels).

**Figure 1.**
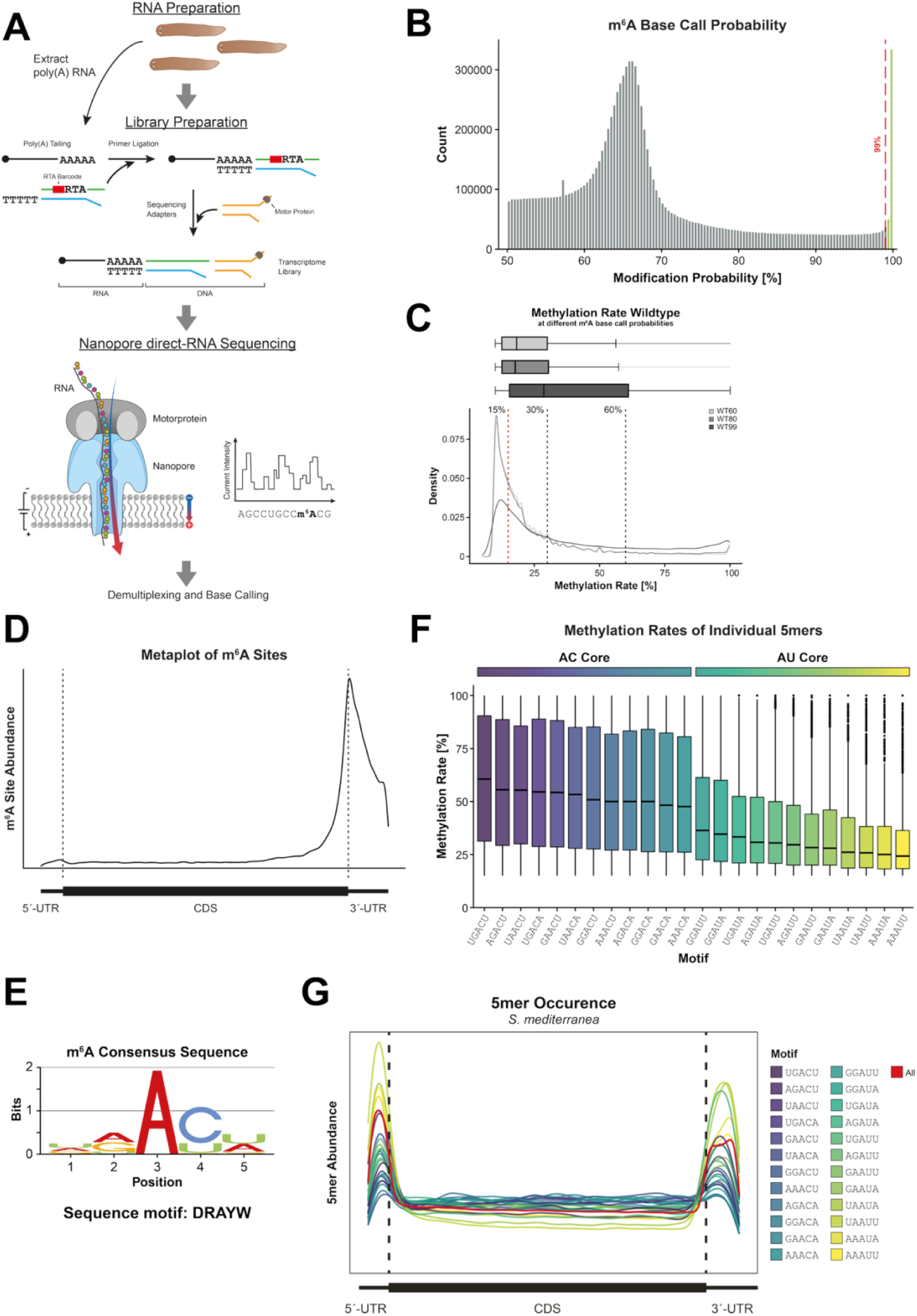
Identification and characterization of high confidence m^6^A sites in *Schmidtea mediterranea*. **(A)** Poly(A) RNA is ligated to a barcoded reverse transcription adapter, followed by attachment of sequencing adapters and a motor protein. Individual RNA molecules are sequenced directly as they translocate through a nanopore, generating characteristic current signals that are used for base calling and demultiplexing, yielding high-quality RNA sequencing data and enabling the detection of RNA modifications. The integration of barcoded adapters allows for multiplexing, which enables high-quality transcriptome profiling with minimized batch effects and detection of m^6^A directly from raw signal (30). **(B)** m^6^A modification probability profile in Nanopore dRNA-Seq reads; the red dashed line represents the applied cutoff (99%), green bars represent the modifications that fulfill this threshold. **(C)** Methylation rates of individual m^6^A sites applying 60%, 80%, and 99% modification probability as cutoffs. Lowering the modification probability threshold increased the fraction of sites with very low apparent methylation rates (≤10%), whereas a 99% cutoff minimized low-confidence calls and was therefore used for high-confidence m^6^A site definition. **(D)** Metagene analysis reveals strong enrichment of m^6^A sites (15% methylation rate in ≥2/3 replicates) at the CDS–3′ UTR junction, with depletion from the 5′ UTR and the CDS. **(E)** Applying a ≥20% contribution threshold at each position within ±2 bp of m^6^A sites yielded a DRAYW consensus motif. **(F)** Individual 5-mer motifs with an AC core exhibit substantially higher methylation rates than AU-core motifs. **(G)** Metagene plots of individual 5-mer consensus motifs reveal preferential enrichment of weakly methylated variants in UTRs, in contrast to the more uniform distribution of highly methylated motifs. Line color represents individual methylation rates as identified in 1F.

We detected a median of 7, 5, and 3 m^6^A sites per planarian gene at methylation rate cutoffs of 15%, 30%, and 60%, respectively (Supplementary Fig. S1A). A metagene analysis of the distribution of these sites across planarian transcripts reveals them to be markedly enriched around stop codons and depleted from the 5’ UTR and CDS (Fig. 1D). This is in line with previous reports (19, 21) and confirms that m^6^A sites in planarians are positioned across transcripts in a manner similar to that observed in humans and other vertebrates (15, 56). In contrast, this pattern differs from that described for *D. melanogaster*, where m^6^A is predominantly localized at the very 5′ ends of transcripts (10).

Next, we used the dRNA-seq data to identify the consensus motif for m^6^A placement in planarians. To this end, we extended called m^6^A sites by ±2 base pairs and extracted the sequences of these 5-mers transcriptome-wide. We considered bases with a contribution of ≥20% at individual positions for the consensus motif. This approach revealed that, in planarians, m^6^A is predominantly installed in a DRAYW sequence context (or DRACW when using 30% and 60% cutoffs for motif methylation rate; with D = not C, R = A or G, Y = C or T and W = A or T) (Fig. 1E). The DRAYW motif in planarians is reminiscent of the consensus motifs previously described for vertebrates (DRACH, with H = not G) (15, 56) and *D. melanogaster* (RRACH) (10). The planarian-specific divergence can likely be attributed to the high AT content of the planarian genome (57), and an adaptation of the writer machinery’s motif recognition from H to W at the +2 position of the m^6^A sequence motif context.

Having identified a consensus motif for m^6^A placement, we next asked which of the individual 5-mers is most efficiently methylated. To that end, we plotted the methylation rate as a function of the methylated 5-mer (Fig. 1F). Interestingly, this analysis revealed the critical importance of a +1 cytosine for efficient methylation by the planarian MTC, as corroborated by GLORI data (21). All 5-mers containing a +1 cytosine (+1C) exhibited average methylation levels exceeding 45%, while the inclusion of a +1 uridine (+1U) resulted in a clear decrease in the average methylation rate to about 25 to 35%. This preference is also evident when comparing motif-specific methylation rates with the occurrence of individual 5-mers (Supplementary Fig. S1B). To confirm that it is solely the interaction of the MTC with the motif that determines the methylation rate and to exclude that the rate is influenced by the location of the individual sites within transcripts, we created a metaplot of individual 5-mer occurrences along planarian transcripts (Fig. 1G). Interestingly, this analysis revealed the significant enrichment of inefficiently methylated motifs in both 5’ and 3’ UTRs. In contrast, strongly methylated motifs are distributed more evenly along transcripts. These observations allow us to draw the following conclusions: First, in the CDS and 3’ UTR of planarian transcripts m^6^A motif occurrence can qualitatively explain the presence of methylated adenosines (Figs. 1D,G). Second, in planarians, unlike in *D. melanogaster* (10, 11), a mechanism must exists that prevents the MTC from accessing the 5’ UTR of transcripts. To corroborate our findings in a better-studied system, we plotted the occurrence of all permutations of the DRACH consensus motif in the human transcriptome (Supplementary Fig. S1C). Intriguingly, and in contrast to the situation in planarians, human transcripts exhibit elevated levels of CDS-based motifs, resulting in a relatively even distribution of DRACH consensus motifs along transcripts. This pattern suggests that, in humans, the MTC needs to be specifically recruited to the STOP codon of transcripts, e.g. by its VIRMA/KIAA1429 subunit (58). Planarians also possess a KIAA1429 homolog (19, 21), which may contribute to MTC recruitment to the stop codon, as suggested by the slight offset between motif occurrence and the peak in m^6^A sites (Figs. 1D,G).

Taken together, multiplexed Nanopore dRNA-seq enabled us to generate a high-confidence atlas of ∼72,200 m^6^A sites across the planarian transcriptome, which - owing to its stringency and robustness - provides a valuable reference resource for the field. Most planarian genes harbor one to five m^6^A sites, strongly enriched near stop codons and depleted from 5′ UTRs and CDS regions. Together with the identification of a planarian-specific DRAYW consensus motif that shows strong dependence on a +1 cytosine for efficient methylation, these findings indicate that planarian m^6^A profiles closely resemble those of vertebrates rather than Drosophila (15, 54, 56).

### Differential methylation frequency of m^6^A motifs at exon starts and ends reveals an altered exclusion zone in planarians

To assess whether the EJC-mediated exclusion zone model of m^6^A placement applies to planarians (7), we analyzed the distribution of m^6^A sites across different length classes of planarian exons. Specifically, we used flexible generalized additive models (GAMs; see Methods) to model a metric that we termed *methylation frequency* - defined as the proportion of consensus DRAYW motifs that are methylated - across a ±500 nt window surrounding splice junctions for internal exon starts, internal exon ends, and last exons (Fig. 2A). Because N^6^-methylated adenosines are directly identified from Nanopore raw signal, we could not omit chemical treatment to derive this metric, as has been done for GLORI data (15, 21). Instead, we defined *methylation frequency* as the ratio of the abundance of methylated consensus DRAYW motifs to the abundance of possible permutations of the motif across the planarian transcriptome (Figs. 1F,G). To examine exon length-dependent trends, we stratified planarian exons into quartiles. For internal exons, these classes comprised exons <242 nt (shortest), 242-415 nt (short), 416-727 nt (long), and >727 nt (longest); for last exons, the corresponding ranges were <259 nt (shortest), 259-401 nt (short), 402-678 nt (long), and >678 nt (longest) (Fig. 2A).

**Figure 2.**
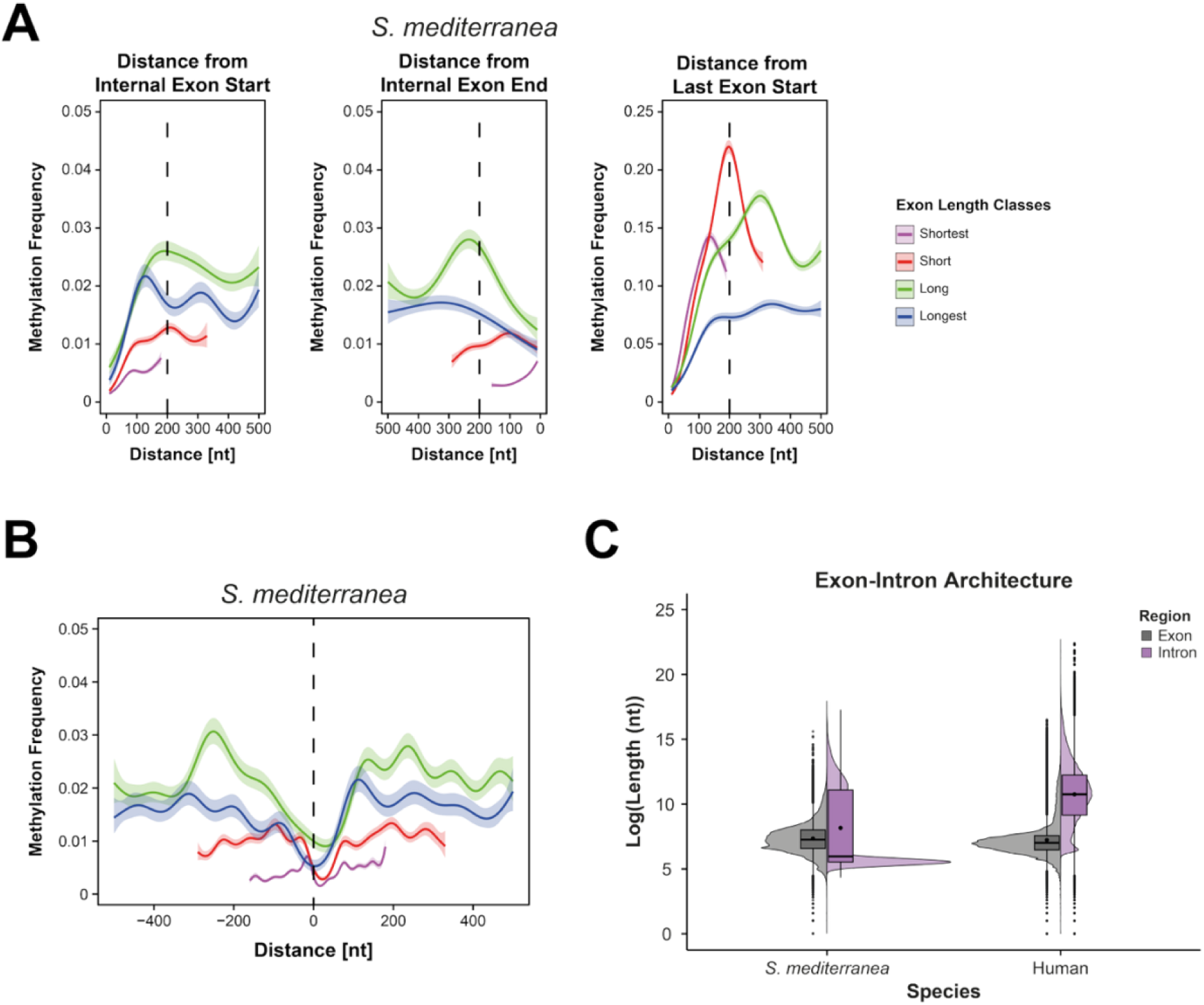
Planarians display an altered m^6^A exclusion zone at exon-exon junctions. **(A)** Generalized additive models (GAMs) of methylation frequency at exon boundaries reveal a pronounced depletion of m^6^A at exon starts, whereas exon ends show no detectable depletion in short exons. GAMs were fitted for exon length classes defined by length quartiles. Shaded areas indicate 95% confidence intervals of the model fit; methylation frequency was tested in bins of 10 nt. **(B)** GAMs across splice junctions uncover a shifted m^6^A exclusion zone ∼20 nt downstream of exon-intron junctions, which is narrowed in short exons and widens with increasing exon length. GAMs were fitted separately for exon length classes defined by length quartiles. Shaded areas denote 95% confidence intervals of the model fit; methylation frequency was tested in bins of 10 nt. **(C)** Comparison of exon-intron architecture highlights major differences between planarian and human genes: planarian genes exhibit a prominent peak of short introns accompanied by a broad distribution of long introns, whereas human genes are characterized predominantly by long introns.

We observed a depletion of m^6^A at the start of both internal and last exons across all length classes (Fig. 2A). In contrast, the extent of m^6^A depletion at internal exon ends varied with exon length. In the shortest exon length class (<241 nt) we did not observe any depletion, however, a shallow exclusion zone up to ∼200 nt from the exon-intron junction became apparent in the two long exon quartiles (Fig. 2A). To obtain a comprehensive view of regions flanking splice junctions, we combined *methylation frequency* profiles from internal exon ends and starts and fitted a continuous generalized additive model across ±500 nt. Apart from the longest exon quartile, m^6^A depletion peaked approximately 20-50 nt downstream of the splice junction in all exon length classes (Fig. 2B). Moreover, with decreasing exon length we found the resulting exclusion zone to become narrower. While its spans ∼100 nt in short exons, it is expanded to 200 nt in long exons, a width comparable to what was observed in humans (Supplementary Figs. S2A,B). This suggests that the EJC might block m^6^A installation near exon ends in longer exons, but not in shorter ones.

To further evaluate the reduced methylation frequency at exon-intron junctions, we repeated the motif occurrence and m^6^A site analyses focusing exclusively on single-exon transcripts. Because these transcripts are not spliced, they do not recruit the EJC and thus provide a direct contrast to multi-exon transcripts, in which EJC deposition could influence m^6^A placement (Fig. 2A). Although motif occurrence did not differ between single-exon and spliced transcripts (Fig. 1G, Supplementary Fig. S2C), m^6^A deposition patterns were clearly distinct. In both transcript classes, 5′ UTR motifs remain largely unmethylated and a prominent 3′ UTR peak dominates the overall m^6^A distribution. However, single-exon transcripts exhibited a notably higher proportion of CDS-localized m^6^A sites (compare Fig. 1D and Supplementary Fig. S2D). This suggests that the EJC likely restricts m^6^A deposition within the CDS of multi-exon transcripts. Nevertheless, because CDS motif density is comparatively low in planarians (Fig. 1G), this effect may be less pronounced than in other species.

In summary, the analysis of CDS-m^6^A methylation patterns in planarians revealed a distribution that is consistent with an EJC-associated exclusion model (7). However, we found the planarian exclusion zone to be dependent on exon length (Figs. 2A,B). Notably, 48.1% of all planarian CDS-linked m^6^A sites reside in exons shorter than 400 nt (methylation rate ≥15%; Supplementary Table S5). These exons are excluded from receiving any methylation under the human-derived EJC exclusion model (7). A proper interpretation of our data thus requires the consideration of planarian gene architecture, which closely resembles that of *Drosophila* and is characterized by a prominent peak of short introns (45-85 bp) alongside a broad distribution of longer introns (59) (Fig. 2C). The short introns likely result in intron-defined splicing, which in turn limits the efficient placement of the EJC in short exons, allowing deposition of m^6^A by the MTC close to exon boundaries (see Discussion).

### EJC depletion partially alters the m^6^A methylome in planarians

To clarify the potential role of the EJC in m^6^A placement in planarians, we next performed RNA interference (RNAi)-induced knockdowns of the planarian EJC components elF4A3 and Y14. Both *eIF4A3* (RNAi) or *y14* (RNAi) resulted in severe phenotypical changes 10 days post first feeding (dpff), which subsequently led to lethality (Supplementary Figs. S3A,B). More specifically, RNAi-treated worms showed massive deformations, accompanied by severely restricted mobility. However, when we used mass spectrometry to measure the influence of *eIF4A3* (RNAi) and *y14* (RNAi) on global m^6^A levels in these animals, we found no changes as compared to wildtype (WT) levels (Supplementary Fig. S3C). Next, we performed Nanopore dRNA-seq at 6 dpff and 8 dpff, following efficient *eIF4A3* (RNAi) and *y14* (RNAi) knockdowns (Supplementary Fig. S3D). Importantly, both time points precede the onset of phenotypic changes, and they both did not lead to significant alterations in global m^6^A levels, as assessed by both mass spectrometry and Nanopore m^6^A base calling (Fig. 3A). If the EJC were to block access to suitable m^6^A motifs close to splice junctions, its loss should lead to increased m^6^A levels in these regions. However, EJC depletion did not significantly change overall m^6^A methylation levels but rather resulted in wild-type-like methylation patterns (Fig. 3B, Supplementary Fig. S3E). Because CDS-linked m^6^A deposition is rare in planarians – as compared to humans - and the exclusion zone is narrow around splice junctions in short planarian exons, we reasoned that any effect that the EJC might have on m^6^A deposition, will be limited to these small areas without causing global changes to the methylation landscape.

**Figure 3.**
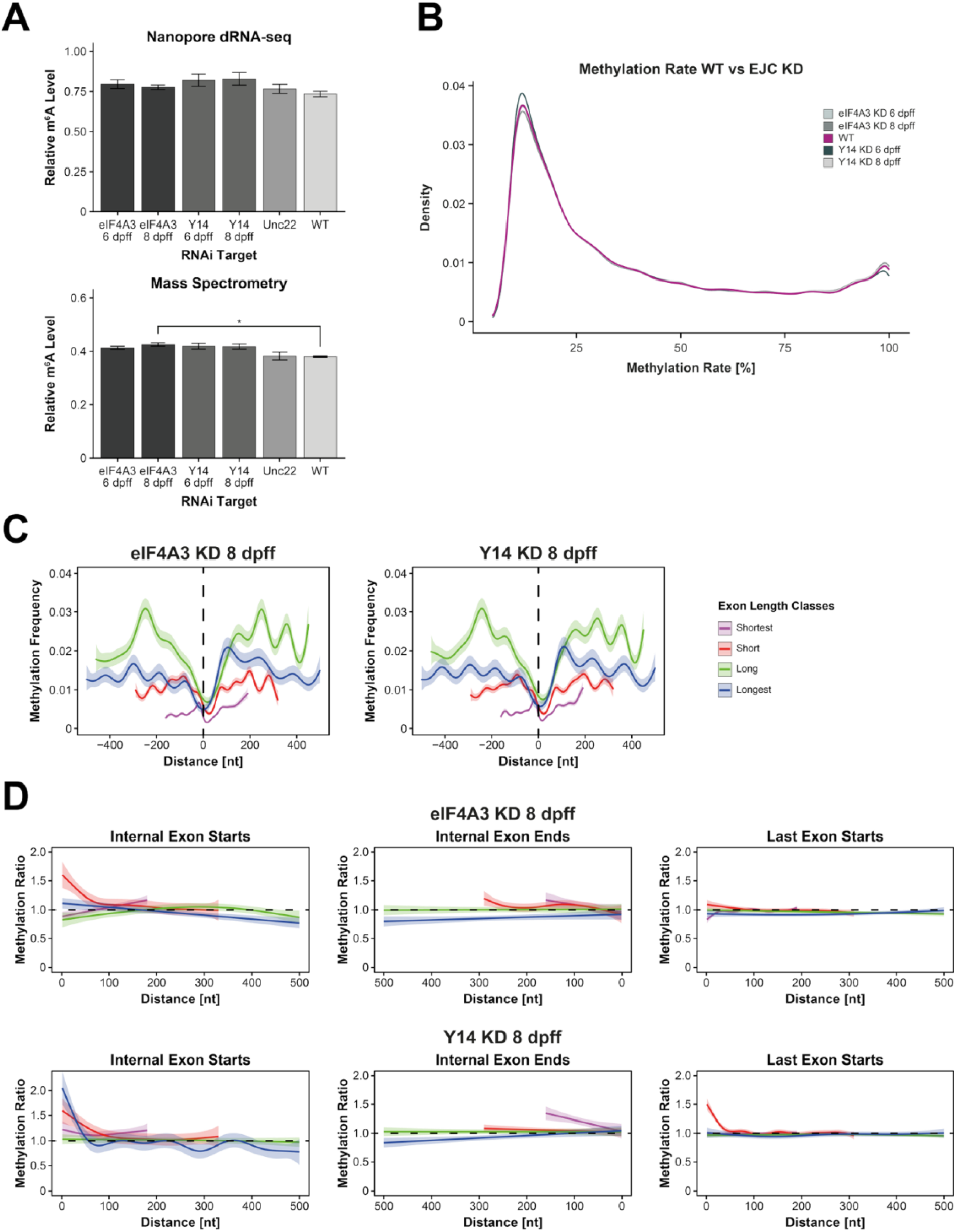
The EJC is responsible for the narrow and shifted exclusion zone in planarians. **(A)** Global m^6^A levels were quantified by mass spectrometry (left) and Nanopore direct RNA sequencing (right) following RNAi-mediated knockdown of the indicated genes. Concordant measurements across both approaches show that EJC depletion does not lead to a global increase in m^6^A levels. Bars represent mean ± SE. Statistical significance was assessed by ANOVA followed by Tukey’s HSD test, with p-values corrected for multiple testing using the Studentized range distribution (adjusted *p* ≤ 0.05). **(B)** Methylation rates of individual m^6^A sites show no marked differences between wildtype and EJC depleted samples. **(C)** Generalized additive models (GAMs) of methylation frequency in *eIF4A3* (RNAi) and *y14* (RNAi) animals across splice junctions at 8 dpff reveal no considerable changes in m^6^A deposition compared to wild type (Fig. 2B). GAMs were fitted for exon length classes, stratified by quartiles of exon length in wild-type (WT) samples. Shaded areas indicate 95% confidence intervals of the model fit; methylation frequency was tested in bins of 10 nt. **(D)** GAMs of methylation frequency ratios between wild type and *eIF4A3* (RNAi) or *y14* (RNAi), respectively, at 8 dpff uncovers a localized increase in m^6^A deposition at exon starts. GAMs were fitted for exon length classes defined by length quartiles in WT. Shaded areas denote 95% confidence intervals of the model fit; methylation frequency ratios were tested in bins of 10 nt.

To assess the impact of EJC depletion on m^6^A deposition near splice junctions, we analyzed methylation frequency as a function of distance following *eIF4A3* (RNAi) and *y14* (RNAi) using generalized additive models (GAMs). At the level of absolute methylation frequency, this analysis did not reveal considerable differences relative to wild type (Fig. 3C, Supplementary Figs. S3F,G, S4A). To increase sensitivity for detecting more subtle, spatially restricted effects, we additionally utilized GAMs to examine ratios of methylation frequency in the knockdown conditions relative to wild type (7). This comparative analysis uncovered an increase in m^6^A methylation frequency toward splice junctions at exon starts in both knockdown conditions, whereas no comparable changes were observed at internal exon ends (Fig. 3D, Supplementary Fig. S4B). Because this increase in *methylation frequency* at exon starts following an EJC knockdown is consistent with our observations in wild type animals (Figs. 2 A,C), our EJC knockdown data confirm that the m^6^A exclusion zone is indeed the result of post-splicing deposition of the EJC on planarian mRNAs. Taken together, we show that the EJC exclusion model near splice junctions (7) is conserved from humans to planarians, albeit with a somewhat shifted exclusion zone dependent on exon length (see Discussion).

### Loss of m^6^A alters gene expression patterns in planarians

To evaluate the impact of m^6^A on transcript fate, we depleted the MTC components METTL3 and METTL14 by RNAi for 24 days, which led to significantly lower transcript levels relative to *unc22* (RNAi) controls (Supplementary Figs. S5A,B). We selected the 24-day time point for our analyses because phenotypic abnormalities – such as lysis spots and head deformation – that culminated in lethality at 41 dpff, only became apparent at 27 dpff (Supplementary Fig. S5C). To confirm that both *mettl3* and *mettl14* (RNAi) knockdowns lead to a significant reduction in m^6^A levels in planarian mRNA, we next analyzed poly(A)-selected RNA from *mettl3* (RNAi) and *mettl14* (RNAi) worms via mass spectrometry. Given that we find no evidence for m^6^A erasers in planarians (Supplementary Table S3), only reduced MTC activity should lead to a reduction in overall m^6^A levels on mRNAs. In line with this assumption, both knockdowns resulted in a substantial depletion of m^6^A from the planarian transcriptome, e.g. relative methylation levels were reduced to 34% in *mettl14* (RNAi) worms and to 44% in *mettl3* (RNAi) animals (Fig. 4A). Because of the more substantial reduction of m^6^A levels following *mettl14* (RNAi), we focused on this knockdown strategy for further analyses and generated dRNA-seq libraries for *mettl14* (RNAi) and *unc22* (RNAi) controls. Whereas *mettl14* (RNAi) libraries averaged ∼4.4M reads (median read length: 631 nt, N50: 936 nt), *unc22* (RNAi) libraries averaged ∼3.9M reads (median read length: 584 nt, N50: 1008 nt) (Supplementary Table S4). Consistent with our mass spectrometry results, m^6^A detection using Oxford’s Dorado software revealed an ∼73.2% reduction in global m^6^A levels in *mettl14* (RNAi) animals (Fig. 4B). Furthermore, methylation at individual sites dropped considerably, with most of the remaining sites showing methylation rates of less than 25% (Fig. 4C). Thus, within the limitations of RNAi-mediated knockdowns in planarians, we achieved a significant reduction of m^6^A in the planarian transcriptome at a time point where phenotypical changes to the animals were not yet detectable. We next defined high confidence m^6^A sites in *mettl14* (RNAi) data as previously described (Supplementary Table S2), using methylation rate cutoffs of 15%, 30%, and 60% in at least two out of three replicates as threshold. In applying these criteria, we observed a significant reduction of m^6^A sites to 20,852 for sites methylated ≥ 15% (down from 72,237 in wildtype samples), to 7,966 for 30% sites (down from 46,520), and to 1,172 for 60% sites (down from 24,968) (Fig. 4D).

**Figure 4.**
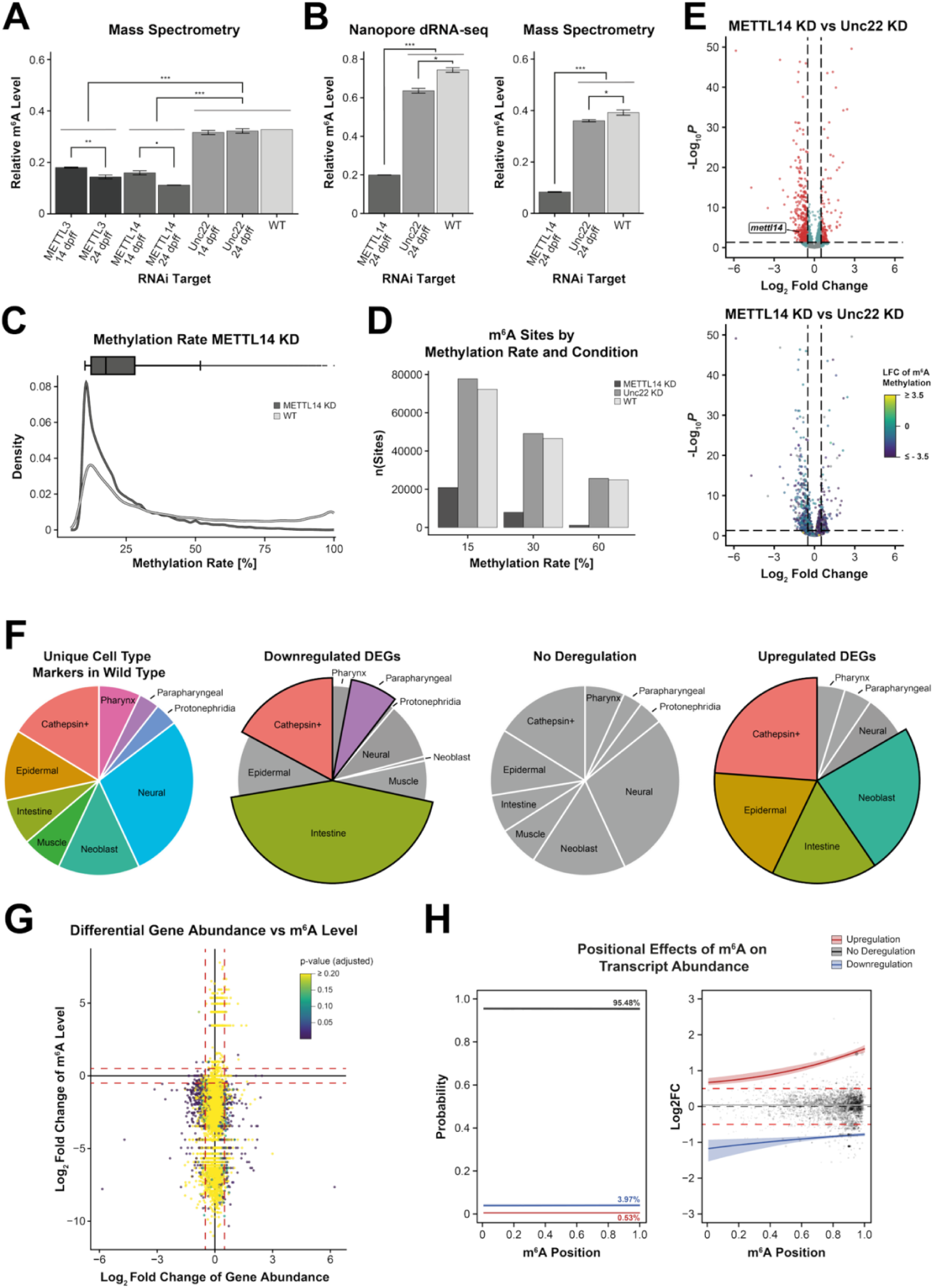
Depletion of the MTC by *mettl14*(RNAi) reveals a significant remodeling of the planarian transcriptome. **(A)** Mass spectrometric quantification of global m^6^A levels following RNAi-mediated knockdown of METTL3 and METTL14. Abundance of m^6^A, normalized to the total of adenine nucleotides (relative m^6^A level), was measured at the indicated time points. Both knockdowns led to a progressive and significant reduction in m^6^A levels compared to controls, with METTL14 depletion showing the strongest effect at 24 dpff, thereby identifying it as the optimal target for subsequent analyses. Bars represent mean ± SE. Statistical significance was assessed by ANOVA followed by Tukey’s HSD test, with p-values corrected for multiple testing using the Studentized range distribution (*** ≤ 0.001; ** ≤ 0.01; * ≤ 0.05; • ≤ 0.1). **(B)** Relative m^6^A levels quantified by mass spectrometry (left) and Nanopore direct RNA sequencing (right) confirm efficient depletion of m^6^A upon knockdown of the MTC core component METTL14. Both approaches consistently demonstrate a significant reduction in global m^6^A levels. Bars represent mean ± SE. Statistical significance was assessed by ANOVA followed by Tukey’s HSD test, with p-values corrected for multiple testing (*** ≤ 0.001; * ≤ 0.05). **(C)** Distribution of methylation rates for individual m⁶A sites in *mettl14* (RNAi) compared to wild type, revealing a pronounced decrease in site-specific methylation rates. **(D)** Number of m^6^A sites identified in wildtype, *mettl14* (RNAi), and *unc22* (RNAi) conditions at methylation rate cutoffs of 15%, 30%, 60%. At all methylation rate cutoffs, *mettl14* (RNAi) consistently showed a substantial reduction in identified m^6^A sites. **(E)** Volcano plot showing gene abundance changes between *mettl14* (RNAi) and *unc22* (RNAi) conditions. Genes with an adjusted p-value ≤ 0.05 and a log_2_ fold change ≤ −0.5 or ≥ 0.5 were classified as significantly deregulated; top: red dots). The corresponding log_2_ fold change in m^6^A site abundance between *mettl14* (RNAi) and *unc22* (RNAi) is indicated by dot color (bottom). **(F)** Distribution of cell-type-specific marker genes across the planarian single-cell atlas. Genes that were not significantly deregulated upon *mettl14* (RNAi) showed no cell-type enrichment. Among significantly downregulated genes, markers for intestine, cathepsin⁺, and parapharyngeal cells were overrepresented, whereas neural, pharyngeal, protonephridial, and neoblast markers were depleted. Conversely, in significantly upregulated genes cathepsin⁺, epidermal, intestine, and neoblast markers were overrepresented, while protonephridial and muscle markers were depleted. **(G)** Scatter plot of log2 fold changes in m^6^A site abundance versus log_2_ fold changes in gene abundance shows no direct relationship between changes in m^6^A and the magnitude or direction of expression changes. Red dashed lines indicate log_2_ fold change thresholds of −0.5 and 0.5; dot color denotes the adjusted p-value for differential expression. **(H)** Bayesian mixture modeling indicates that m^6^A is more likely to be associated with transcript downregulation than upregulation, independent of its position along the transcript. Similarly, the magnitude of deregulation is largely position independent. The unregulated transcript class (gray line) shows the highest probability and centers around log₂ fold changes of ∼0. Shaded areas represent 80% credibility intervals.

Next, we investigated the impact of m^6^A depletion on the planarian transcriptome. To achieve this, we removed the few rRNA and tRNA reads present, mapped the remaining reads to the planarian transcriptome, and quantified gene expression using Salmon (41). We then employed DESeq2 (43) to investigate differential gene expression between *mettl14* (RNAi) and *unc22* (RNAi) following m^6^A depletion (Fig. 4E). Importantly, this approach minimizes confounding RNAi-induced effects, which were apparent when comparing *mettl14* (RNAi) vs. WT and *unc22* (RNAi) vs. WT (Supplementary Fig. S5D). In total, 1,059 genes are significantly deregulated upon *mettl14* (RNAi), including 413 upregulated and 645 downregulated genes (Fig. 4E, Supplementary Table S6). Given previous reports of altered cell type composition in *S. mediterranea* following *kiaa1429* (RNAi), a component of the MTC (19), we asked whether specific cell types were preferentially affected in our data set. To that end we leveraged expression data from the planarian single-cell atlas (45) and extracted genes uniquely associated with individual cell types to generate a reference set of cell-type-specific markers (Methods). Mapping *mettl14* (RNAi)-deregulated genes onto this marker set revealed that upregulated genes were enriched for markers of cathepsin⁺, epidermal and intestine cells, as well as for neoblasts, while being depleted for markers of protonephridia and muscle cells, which were not represented at all (Fig. 4F). In contrast, downregulated genes were dominated by intestine cell markers and further showed enrichment for cathepsin⁺ and parapharyngeal markers, alongside the depletion of neural, pharyngeal, protonephridial, and neoblast markers, with the latter two cell types being almost completely absent. Notably, genes that were not significantly deregulated exhibited no bias toward any cell-type-specific marker set, highlighting the specificity of cell type identity patterns observed for deregulated genes. These findings are consistent with previous studies showing that disruption of the m^6^A pathway primarily impairs lineage differentiation rather than global stem cell maintenance (18, 19, 21). Loss of intestine cells due to depletion of intestine progenitors, accompanied by accumulation of neural-like progenitors, has been reported following m^6^A depletion (19). Defects in protonephridia and muscle lineages, as well as impaired neural regeneration, were also observed upon m^6^A loss (18). Similarly, knockdown of YTHDF family m^6^A readers predominantly affected differentiation of parenchymal, cathepsin⁺, and neural lineages while leaving the neoblast pool largely intact (21). Together, these studies support a model in which m^6^A regulates lineage-specific differentiation programs, consistent with the cell-type-dependent stability effects identified here.

Notably, 90.5% of all deregulated genes (958 of 1,059) carried m^6^A marks, suggesting that altered m^6^A status is the major driver of the observed expression changes. To explore this further, we analyzed log_2_ fold changes of m^6^A sites with methylation rates ≥15%. This revealed no direct correlation between loss or gain of m^6^A sites and either the direction or magnitude of gene expression changes (Fig. 4G). Furthermore, these results indicate that m^6^A broadly influences the fate of modified transcripts but does not act uniformly to stabilize or destabilize them, likely reflecting context-dependent effects mediated by distinct m^6^A reader proteins. Last, prompted by the skewed distribution of m^6^A sites along planarian transcripts, with strong enrichment near the stop codon (Fig. 1D), we hypothesized that the position of an m^6^A mark within a transcript modulates its effect on transcript fate. To test this, we employed a Bayesian mixture model to assess whether the average position of m^6^A sites influences probability, direction and magnitude of the log_2_ fold change of transcript abundance following *mettl14* (RNAi) (Methods). For this analysis, only transcripts losing ≥80% of their m^6^A sites were considered (n = 3,966), with m^6^A site loss being defined as the reduction in methylation rate per sites from ≥15% to 0% following *mettl14* (RNAi) (Supplementary Fig. S5E). Consistent with previous observations (Figs. 4E,G), we found m^6^A to be associated with both stabilizing and destabilizing effects on transcripts. However, the model indicates that most transcripts (∼95.5%; 80% credibility interval: 94.82–96.10%) exhibit no substantial change in abundance following *mettl14* (RNAi) (Fig. 4H). Downregulation has a probability of 3.97% (80% credibility interval: 3.39–4.62%). Importantly, for these downregulated transcripts, the magnitude of the log₂ fold change did not substantially vary with the mean position of m^6^A sites along the transcript. In contrast, upregulation was rare (probability: 0.53%; 0.38–0.70%). While the model captures the overall distribution of effects, particularly on the downregulated side, posterior predictive checks indicate an overestimation of strong positive effects (>0.5) of the model. Accordingly, conclusions regarding the probability and magnitude of such effects should be interpreted with caution, and that such effects may even be less likely than indicated by the model. Overall, the results suggest that m^6^A more often stabilizes than destabilizes planarian transcripts. Importantly, the magnitude of downregulation is largely independent of m^6^A position, which argues against a functionally enhanced significance of the frequently occurring sites near the stop codons of transcripts (Fig. 1G).

### Poly(A) tail shortening in a METTL14 knockdown occurs independently of m6A status

Nanopore dRNA-Seq offers the unique possibility to directly measure poly(A) tail length. This benefits our study as both m^6^A sites and poly(A) tail length are involved in pre-mRNA surveillance. Moreover, prior evidence has begun to connect the m^6^A machinery to cleavage and (alternative) polyadenylation (58, 60, 61). When we analyzed the length of poly(A) tails in our data, we found that they were significantly shorter in *mettl14* (RNAi) animals (mean: 75 nt) as compared to WT and *unc22* (RNAi) animals (mean: 85 nt and 87 nt respectively) (Fig. 5A). To assess whether this effect was caused by m^6^A, we classified the planarian wildtype transcriptome into two groups: transcripts that harbor m^6^A sites and transcripts that do not. In all conditions analyzed (wildtype, *unc22* (RNAi), and *mettl14* (RNAi)), m^6^A-marked transcripts exhibited considerably longer poly(A) tails compared to non-m^6^A transcripts (Fig. 5B). However, we observed that poly(A) tails were uniformly shorter in *mettl14* (RNAi) animals, regardless of their m^6^A status in wildtype conditions. To further examine a potential relationship between m^6^A and poly(A) tail length, we grouped genes according to previously calculated m^6^A log₂ fold changes between *mettl14* (RNAi) and *unc22* (RNAi) conditions. Genes were assigned into four categories: transcripts that lose m^6^A sites (LFC < −0.5), transcripts that gain m^6^A sites (LFC > 0.5), transcripts with unchanged m^6^A levels (−0.5 ≤ LFC ≤ 0.5), and transcripts that do not carry detectable m^6^A. Surprisingly, we observed poly(A) tail shortening across all gene classes upon *mettl14* (RNAi), including for transcripts that lack m^6^A (Fig. 5C). This finding was unexpected, as m^6^A-marked transcripts display longer poly(A) tails under wild-type conditions (Fig. 5B). Thus, together, these results suggest that a METTL14 knockdown broadly disrupts poly(A) tail homeostasis and mRNA surveillance, likely through both direct effects on m^6^A deposition and indirect effects on global poly(A) tail regulation (see Discussion).

**Figure 5.**
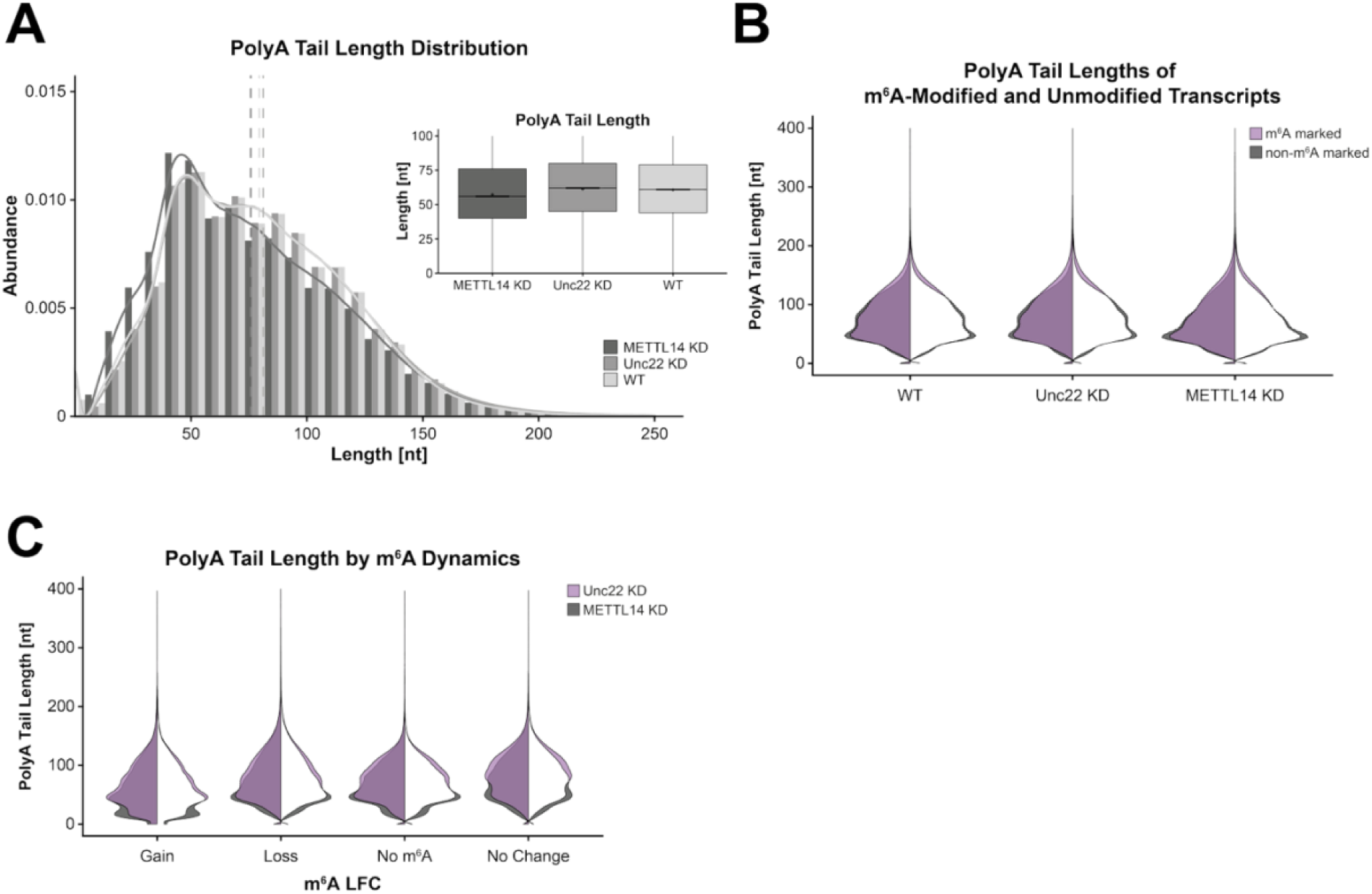
A METTL14 Knockdown leads to globally shortened poly(A) tails. **(A)** Distribution of poly(A) tail length in wildtype, *mettl14* RNAi), and *unc22* (RNAi) conditions. dRNA-Seq reads of *mettl14* (RNAi) displayed significantly shorter poly(A) tails than those of *unc22* (RNAi) or wildtype (adjusted p-value = 0). **(B)** Poly(A) tail length in wildtype, *mettl14* (RNAi), and *unc22* (RNAi) stratified by m^6^A status in wildtype. Split violin plots display the poly(A) tail length distribution of transcript classes; left: length distributions of both classes, right: differences in length distributions (lilac = poly(A) tail length distribution of m^6^A-marked transcripts (WT); grey = poly(A) tail length distribution of transcripts without m^6^A detected (WT)). m^6^A transcripts had longer poly(A) tails than transcripts without m^6^A. Poly(A) tail length in *mettl14* (RNAi) conditions were shorter irrespective of m^6^A status. **(C)** Genes were clustered by m^6^A log_2_ fold change (m^6^A loss: LFC < -0.5; m^6^A gain: LFC > 0.5, No m^6^A: no methylation in either condition; No Change: LFC **≥** -0.5 & **≤** 0.5). Split violin plots display the poly(A) tail length distribution in *mettl14* (RNAi) and *unc22* (RNAi) conditions; left half: length distributions in both conditions, right half: differences in length distributions (lilac = poly(A) tail length distribution in *unc22* (RNAi); grey = poly(A) tail length distribution in *mettl14* (RNAi). All four clusters displayed shorter poly(A) tails in *metll14* (RNAi) than in *unc22* (RNAi).

### m^6^A has opposing effects on transcript decay depending on cellular context

Our previous observations revealed pronounced changes in gene expression as well as a global reduction in poly(A) tail length upon m^6^A depletion (Figs. 4E,F and 5A,B), strongly implicating m^6^A in being involved in the regulation of mRNA stability in planarians. These findings furthermore suggest that altered transcript decay dynamics may underlie a substantial fraction of the observed expression changes. To directly test this hypothesis and to quantify the impact of m^6^A on transcript stability, we next employed an Actinomycin D assay to monitor mRNA decay kinetics over time. Actinomycin D is a well-established transcriptional inhibitor that enables measurement of mRNA stability by acutely blocking transcription and allowing decay of pre-existing transcripts to be monitored over time (62). This approach has previously been used to demonstrate the requirement for ongoing transcription during regeneration, supporting the applicability of this drug in planarians (63). In more detail, we repeated the *mettl14* (RNAi) knockdown as previously described (Supplementary Fig. S5A and Methods) and treated worms with Actinomycin D at 22 dpff. RNA from wildtype, *unc22* (RNAi) and *mettl14* (RNAi) animals was collected 0, 6 and 12 h after this treatment, ribo-depleted and subjected to Illumina paired-end sequencing (Methods). Importantly, to minimize potential biases in mRNA enrichment caused by shortened poly(A) tails (Figs. 5A,B), we used ribo-depleted RNA rather than poly(A)-selected RNA for this analysis. To identify genes displaying differential abundance trajectories during transcriptional arrest, the resulting data were then subjected to a time-series analysis with maSigPro (53). To focus on m^6^A-dependent effects on transcript stability, we restricted this analysis to transcripts harboring at least one m^6^A site with a methylation rate ≥15% under wildtype conditions (based on our prior methylome analysis, Supplementary Fig. S1A, Supplementary Table S2). To reduce noise, we excluded genes whose transcript abundance increased over time, likely reflecting incomplete Actinomycin D penetrance into the entire animal or the activation of compensatory pathways such as drug efflux mechanisms (64). In more detail, gene-wise linear models were fitted separately for each condition (wildtype, *unc22* (RNAi), or *mettl14* (RNAi)), and genes with a significantly positive slope and a log2 fold change ≥0.2 in any condition were removed from the analysis. Subsequently, all remaining genes were centered to their respective 0 h time points and reanalyzed using maSigPro. Following filtering, quadratic regression identified 2,989 genes whose abundance profiles differed significantly between conditions over time (FDR = 0.0027). Subsequent stepwise regression (“two-way forward” method) refined this set to 503 genes that showed significant differences in decay behavior in *mettl14* (RNAi) animals as compared to both wildtype and *unc22* (RNAi) control conditions. To further classify these genes based on their decay dynamics, we determined k = 2 as the optimal number of clusters using principal component analysis and within-cluster sum-of-squares metrics (Fig. 6A, Supplementary Figs. S6A,B and Methods). In the first cluster (n = 187), genes displayed significantly faster transcript decay upon *mettl14* (RNAi), indicating that m^6^A might serve to stabilize these mRNAs. In contrast, genes in the second cluster (n = 316) exhibited slower decay rates in *mettl14* (RNAi) as compared to wild type and *unc22* (RNAi) animals, suggesting that m^6^A promotes transcript decay for this subset (Supplementary Table S7).

**Figure 6.**
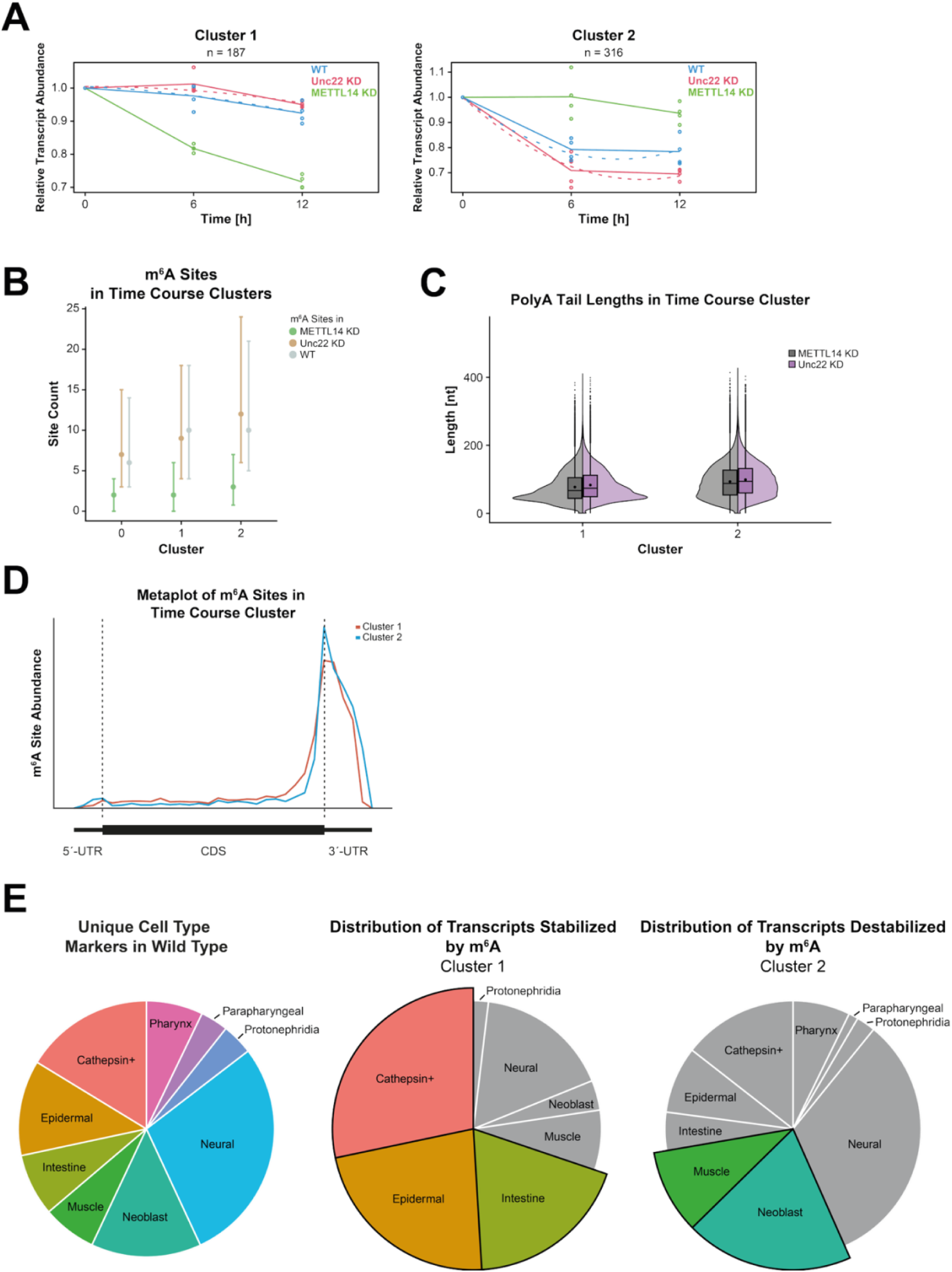
Time series analysis of Actinomycin D assay reveals distinct mRNA stability clusters. **(A)** Time series analysis of transcript stability following *mettl14* (RNAi) identified two distinct gene clusters with opposing decay behaviors. Cluster 1 transcripts (n = 187) exhibited accelerated decay upon m^6^A loss, indicating a stabilizing role of m^6^A, whereas Cluster 2 transcripts (n = 316) showed increased stability, consistent with m^6^A-mediated destabilization. Solid lines represent the mean expression values, while dashed lines indicate the regression fit curves generated by the model. **(B)** Genes assigned to either stability cluster harbored significantly more m^6^A sites than non-clustered genes in both *unc22* (RNAi) and wildtype conditions (adjusted p-values = 4.752 × 10⁻² / 9.392 × 10⁻^4^ for Cluster 1; 1.065 × 10⁻¹³ / 6.797 × 10⁻¹¹ for Cluster 2, respectively). In *unc22* (RNAi) animals, Cluster 2 genes additionally had significantly more m^6^A sites than Cluster 1 genes (padj = 9.598 × 10⁻^4^). Dots indicate medians and whiskers the first and third quartiles. Statistical significance was assessed using the Kruskal–Wallis test with Bonferroni correction. **(C)** Poly(A) tail length was reduced in both clusters upon *mettl14* (RNAi), indicating that differential stability outcomes are not explained by opposing poly(A) tail dynamics. **(D)** Metagene analysis of m^6^A site distribution revealed persistent enrichment toward the CDS–3′UTR junction in both clusters. Notably, Cluster 1 genes displayed slightly higher m^6^A density within the CDS compared to Cluster 2 genes. **(E)** Cell-type marker analysis using the planarian single-cell atlas (45) showed distinct lineage associations for each cluster. In Cluster 1 intestine, epidermal, and cathepsin⁺ markers were overrepresented and neoblast, neural, pharyngeal, and parapharyngeal markers were depleted. In contrast, muscle, and neoblast markers were overrepresented, while protonephridia, intestine, and parapharyngeal markers were depleted in Cluster 2.

In building on these findings, we sought to identify determinants underlying the observed gene-specific differences in mRNA decay. We first examined the m^6^A status of the clustered genes and found that they carried significantly more m^6^A marks than non-clustered genes, supporting that the observed stability effects are m^6^A-dependent (Fig. 6B). Notably, genes in cluster 2, in which m^6^A promotes transcript destabilization, displayed significantly higher overall methylation levels than genes in cluster 1, which we found to be stabilized by m^6^A (Fig. 6B). Moreover, genes within stability clusters exhibited no significant difference in log₂ fold change of m^6^A site abundance compared to non-clustered genes. This observation underscores that the influence of m^6^A on transcript stability is mediated by gene-specific regulatory mechanisms, rather than a uniform response to overall m^6^A depletion (Supplementary Fig. S6C). We next assessed whether differences in poly(A) tail length could account for the opposing decay behaviors. Poly(A) tail analysis revealed a general shortening of poly(A) tails in *mettl14* (RNAi) animals compared to *unc22* (RNAi) controls, irrespective of cluster assignment (Fig. 6C). This indicates that differential mRNA stability between clusters is not primarily driven by poly(A) tail modulation, but likely reflect alternative mechanisms, such as context-dependent recognition of m^6^A sites by distinct m^6^A reader proteins (65–68) (see Discussion). Finally, we examined the positional distribution of m^6^A marks within transcripts. In both clusters, m^6^A sites were predominantly enriched in the vicinity of the stop codon, in line with the metagene plot for overall m^6^A distribution in planarian transcripts (Fig. 1D). However, transcripts in cluster 1, which are stabilized by m^6^A, exhibited modestly higher m^6^A levels within the CDS compared to transcripts in cluster 2 (Fig. 6D). This pattern contrasts with observations in human cells, where CDS-localized m^6^A sites were reported to promote transcript destabilization via translation-dependent decay (9). Taken together with the very low overall frequency of m^6^A sites within the CDS of planarian transcripts (Fig. 1D), our data thus argue against a major role for translation-dependent m^6^A-mediated mRNA decay in planarians.

As a final step in our analyses, we asked whether the effect of m^6^A on transcript stability depends on the cellular context. To address this, we leveraged the planarian single-cell atlas (45) and, as described previously (Fig. 4F and Methods), mapped genes from the identified clusters onto a reference set of cell-type-specific marker genes to assess cell-type enrichment (Fig. 6E). Genes from cluster 1, in which m^6^A stabilizes transcripts, were overrepresented in markers of cathepsin⁺, epidermal, and intestine cells, while being depleted among markers of neoblasts, neural, pharyngeal, and parapharyngeal cells, with the latter two cell types being completely absent from this cluster. In contrast, genes from cluster 2, in which m^6^A promotes transcript decay, were significantly enriched for muscle and neoblast markers but depleted for markers of protonephridia, intestine, and parapharyngeal cells. Notably, these enrichment patterns are largely reciprocal between the two clusters, indicating that m^6^A-mediated transcript stabilization and destabilization preferentially occur in distinct cellular contexts rather than affecting the same cell types in opposing ways. More specifically, our Actinomycin D experiments suggest that m^6^A predominantly exerts stabilizing effects in cathepsin⁺, epidermal, and intestine cells, whereas it preferentially promotes transcript decay in neoblasts and muscle cells. Notably, previous studies have reported that depletion of the m^6^A writer complex primarily affects intestine, muscle, protonephridia, neural, and specific neoblast subpopulations (18, 19), consistent with our observations. Collectively, our results indicate that the impact of m^6^A on mRNA stability is strongly cell-type dependent and likely reflects differences in the expression or activity of m^6^A reader proteins, rather than intrinsic properties of the modified transcripts themselves.

## DISCUSSION

### Principles of m^6^A deposition in the planarian transcriptome

In this study, we delineate the m^6^A methylation landscape of the planarian *Schmidtea mediterranea* and uncover both conserved and cell type-specific features of m^6^A regulation. Leveraging direct RNA sequencing, we quantified absolute m^6^A levels, mapped modification sites at single-nucleotide resolution, and assessed transcript abundance and poly(A) tail dynamics in a single experiment. We show that the core components of the MTC are conserved from humans to planarians and that m^6^A is a prevalent internal modification of planarian mRNAs. Notably, we find no evidence for m^6^A demethylases, indicating that m^6^A marks are not actively removed once deposited. This suggests a comparatively stable methylation landscape in which regulatory specificity is likely primarily achieved through controlled deposition and context-dependent reader engagement rather than dynamic erasure, a model further supported by our stability analyses. Planarian m^6^A sites conform to a DRAYW consensus motif closely related to vertebrate (and even insect) motifs, with modest deviations likely reflecting the high AT content of the planarian genome. At the transcript level, m^6^A is strongly enriched near stop codons and within 3′ UTRs, closely resembling vertebrate profiles and contrasting with the pronounced 5′ bias observed in *Drosophila*.

Despite this overall conservation, m^6^A placement relative to splice junctions confirms that the canonical EJC exclusion model (7) operates in planarians, albeit with an exclusion zone shifted downstream of the splice junction and dependent on exon length. While m^6^A deposition is consistently depleted at exon starts in an EJC-dependent manner, its effect at exon ends varies with exon length: in shorter exons, we observe no exclusion zone at exon ends, while in longer exons, the exclusion zone widens to ∼ 200 nt from the junction. Previous studies have shown that the exon-intron architecture in *Drosophila* is dominated by relatively short introns (< 100 nt), resulting in intron-defined splicing (69, 70). In intron-defined splicing, splice sites are recognized within individual introns (e.g., U1 snRNP binding to the 5′ splice site and U2AF binding to the 3′ splice site of the same intron). This contrasts with exon-defined splicing in vertebrates, where splice sites are recognized across exons (71). An intron-defined splicing reaction (e.g. in *Drosophila*) results in rapid splicing kinetics, which in turn does not allow for efficient EJC positioning *post* splicing and thus results in inefficient nonsense-mediated decay (72, 73). In contrast, in exon-defined splicing, the EJC is typically deposited ∼20–24 nt upstream of exon-exon junctions (74). Because our analysis of planarian exon-intron architecture demonstrates that it closely resembles the situation in *Drosophila* (Fig. 2C), we hypothesize that intron-defined splicing dominates in planarians, at least for the large numbers of small introns. This in turn leads to inefficient deposition of the EJC, which results in the observed dependence of the EJC-mediated exclusion zone on exon length. To determine the precise position of the EJC on planarian transcripts further experiments, such as CLIP (crosslinking and immunoprecipitation) on the planarian EJC, which comprised all components found in vertebrates, are needed.

In contrast to vertebrate systems, where dynamic m^6^A erasure enables rapid transcriptomic remodeling (75), planarians may achieve regulatory specificity of m^6^A placement primarily through elevated mRNA turnover - e.g. high intracellular RNase activity (76) -, and cell-type-specific m^6^A reader activity (see separate Discussion paragraph and (21)). Such regulatory strategy is well suited to planarian biology, where cell division is restricted to neoblasts and regeneration depends on the robust and sustained execution of lineage-specific gene expression programs. The conservation of a vertebrate-like m^6^A distribution in an invertebrate species further suggests strong evolutionary pressure to maintain post-transcriptional control mechanisms that support complex tissue organization and cellular plasticity. Future comparative analyses of m^6^A motifs and reader proteins across regenerative and non-regenerative flatworms may help identify features linked to regenerative capacity. Moreover, integrating single-cell transcriptomics with m^6^A mapping approaches (77) could further resolve how cell-type-specific m^6^A deposition and reader engagement shape gene regulatory programs during regeneration.

### Context matters: m^6^A directs lineage-specific mRNA decay

Furthermore, our findings provide a mechanistic framework that reconciles previously reported phenotypes of m^6^A writer complex depletion with the cell-type-specific effects observed here (18, 19, 21). Time-resolved Actinomycin D assays following *mettl14*(RNAi) identified two distinct gene clusters with opposing stability responses to m^6^A loss: one in which m^6^A stabilizes transcripts and another in which it promotes decay. Importantly, these effects are not uniformly distributed across the transcriptome but are strongly cell-type dependent. Transcripts stabilized by m^6^A are enriched in intestine, epidermal, and cathepsin⁺ cells, whereas transcripts whose decay is promoted by m^6^A are preferentially associated with neoblasts and muscle cells (Fig. 6E). This context-dependent regulation provides a compelling explanation for prior observations that knockdown of m^6^A writer components such as *kiaa1429* or *wtap* does not ablate the general neoblast pool but instead leads to the accumulation of X2 progenitors - particularly neural progenitor-like cells - alongside depletion of intestine and muscle progenitors and their differentiated descendants (18, 19). Our data suggest that loss of m^6^A destabilizes lineage-specific gene programs in intestine, epidermal and cathepsin⁺ cells, impairing differentiation and maintenance of these tissues. Conversely, in neoblasts, where m^6^A normally promotes transcript turnover, its depletion leads to aberrant stabilization of progenitor-associated transcripts, potentially trapping cells in an undifferentiated or lineage-biased progenitor state. This model explains how neoblast identity can remain largely intact while differentiation trajectories are selectively disrupted. Notably, poly(A) tail shortening upon m^6^A depletion occurred across both gene clusters, indicating that differential transcript stability is not primarily driven by poly(A) tail modulation. Instead, our results point to m^6^A reader proteins as key mediators of the opposing stability outcomes. We speculate that lineage-specific expression or activity of m^6^A readers underlies the contrasting effects of m^6^A on transcript fate across cell types.

### m^6^A exerts its opposing effects on mRNA stability independently of poly(A) tail length

Our poly(A) tail analyses (Fig. 5) indicate that m^6^A might contribute to mRNA surveillance through both direct and indirect mechanisms. Directly, loss of m^6^A from transcripts that normally carry the modification abolishes the stabilizing influence of m^6^A-bound readers, leading to their accelerated deadenylation or their impaired polyadenylation. This interpretation is supported by the longer poly(A) tails observed on m^6^A-marked transcripts. These results are also consistent with emerging evidence linking m^6^A to protection of poly(A) tails and regulation of deadenylation dynamics (60, 61). Indirectly, however, METTL14 depletion causes a global shortening of poly(A) tails, including in transcripts that do not normally harbor m^6^A. This unexpected effect suggests that disruption of the MTC is likely to broadly perturb mRNA surveillance pathways. Alternatively, a METTL14 knockdown may elicit a stress response that globally accelerates deadenylation independently of the m^6^A status of specific transcripts. The observation that even transcripts classified as m^6^A-gaining (LFC > 0.5) exhibit shortened poly(A) tails further supports the idea that compensatory or ectopic methylation under METTL14 depletion is insufficient to restore reader-mediated stabilization (Fig. 5C). Together, these findings indicate that intact m^6^A writer activity is required globally for proper poly(A) tail homeostasis.

### Cell-type-specific m^6^A readers dictate divergent mRNA stability programs

The opposing effects of m^6^A on transcript stability across planarian cell types are likely mediated by selective recruitment of stabilizing or decay-promoting machinery by distinct m^6^A readers. A recent characterization of the expanded *ythdf* family in planarians revealed pronounced tissue-specific expression patterns, with *ythdf*-A enriched in the intestine, ythdf-B in the brain, and *ythdf*-C in the lining of the intestine and brain, alongside broader but overlapping expression domains (21). Strikingly, these patterns align closely with our Actinomycin D stability data: m^6^A stabilizes transcripts in epidermal, intestine, and cathepsin⁺ cells with the latter two tissues enriched for *ythdf*-A (*ythdf*-A, -B, and -C were found to be enriched in the epidermis) - whereas cell types enriched for *ythdf*-B and *ythdf*-C, including neural, muscle, and pharyngeal cells, are overrepresented among genes for which m^6^A promotes transcript decay. This correlation supports a model in which *ythdf*-A predominantly confers transcript stabilization, while *ythdf*-B and *ythdf*-C favor decay, reminiscent of mammalian systems in which YTHDF proteins recruit distinct effector complexes to regulate mRNA fate. Consistent with this view, mammalian YTHDFs promote mRNA decay via the CCR4–NOT complex (78), whereas IGF2BP proteins stabilize transcripts by protecting them from deadenylation or by enhancing their translation (79). Notably, we identified additional planarian homologs of m^6^A readers, including IGF2BP, YTHDC1, and HNRNP family members, raising the possibility that stabilizing effects observed in intestine and cathepsin⁺ cells may involve readers beyond the YTHDF family (Supplementary Table S3). At the same time, the broad co-expression of *ythdf* genes suggests potential functional redundancy or combinatorial regulation. Together, these observations indicate that m^6^A-dependent control of transcript stability in planarians might be dictated primarily by the local reader landscape, underscoring the need for future studies to dissect reader-specific functions, redundancy, and downstream effector pathways in a cell-type-resolved manner.

## ACKNOWLEDGEMENTS

We thank Miklas Hafke, Jule Lengsfeld, and Simone Keß for conducting preliminary experiments. We thank Wiep van der Toorn for advice and support with demultiplexing Nanopore libraries using WarpDemuX. We thank Catherine Broberg Vågbø from the Proteomics and Modomics Experimental Core Facility (PROMEC) of the Norwegian University of Science and Technology (NTNU) for carrying out mass spectrometry measurements and advice in result evaluation. We thank Vladyslava Gorbovytska for advice regarding the graphical abstract and for critically reading the manuscript.

## AUTHOR CONTRIBUTIONS

C.H., A.P. and C.-D.K. conceived and designed all experiments. C.H. carried out all experiments, except for dRNA-seq library preparations. A.P. performed all computational data analysis with advice from R.P.S. on dRNA-seq data analysis, except for the Bayesian hierarchical mixture-of-experts model. A.-S.G.-B. performed all dRNA-seq library preparation. Nanopore sequencing was carried out by R.P.S. and A.-S.G.-B. GAM analysis was established by L.H. and subsequently carried out by A.P. L.H. developed and applied the Bayesian hierarchical mixture-of-experts model. A.P., C.-D.K., and C.H. drafted the manuscript with input from all authors.

## SUPPLEMENTARY DATA

Supplementary Data (Supplementary Figures S1-S6 and Supplementary Tables S1-S7) are available with this article under (*link to be inserted here*).

## DATA AVAILABILITY

All sequencing data (Nanopore and short read) generated in this study have been deposited in the NCBI Gene Expression Omnibus (GEO) database *(accession code pending)*.

## CODE AVAILABILITY

All code for the Bayesian Mixture model analysis is available at GitHub under https://github.com/LisaHuelsmann/m6ABayesianMixture.

