## Supplementary_Material for "mRNA stability in response to m^6^A placement is linked to cell identity in planarians"

List of supplementary material:

Supplementary Figures S1 – S6

Supplementary Tables S1 – S7

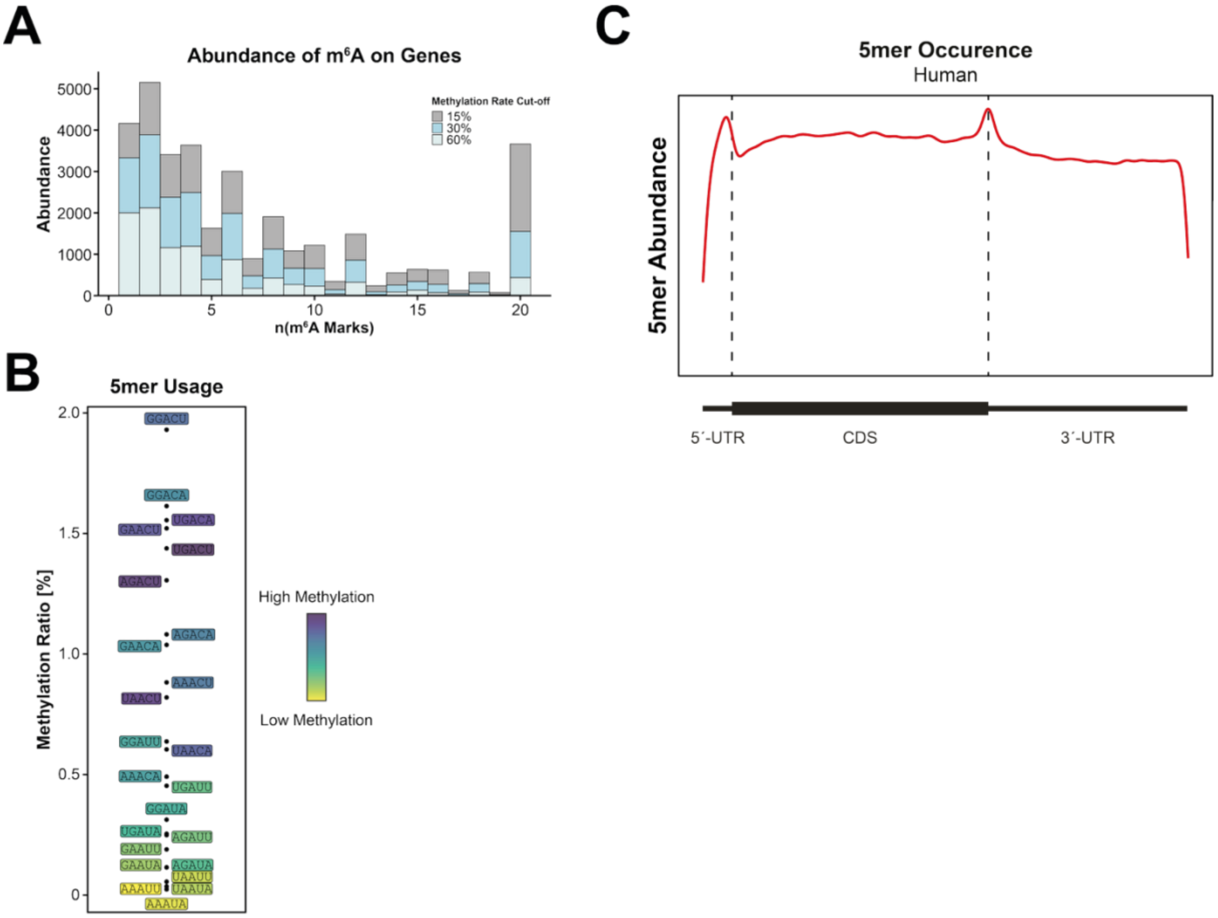


**Figure S1. m$^6$A site occurrence in planarians. (A)** Number of m$^6$A sites on planarian genes at different methylation rates. Median number of sites per gene was 7 at 15%, 4 at 30%, and 4 at 60% methylation rate, respectively. **(B)** Methylation ratio of individual 5-mers (defined as methylated 5-mer/occurrence of 5-mer) coincides with methylation rate of respective 5-mers, meaning that 5-mers with lower methylation rates are also methylated less frequently in total. Displayed colors are equivalent to Fig. 1F. **(C)** Metagene plot of all 5-mers making up the human DRACH motif shows that motifs are distributed more evenly along human genes with less pronounced enrichment around the CDS-3'UTR border.

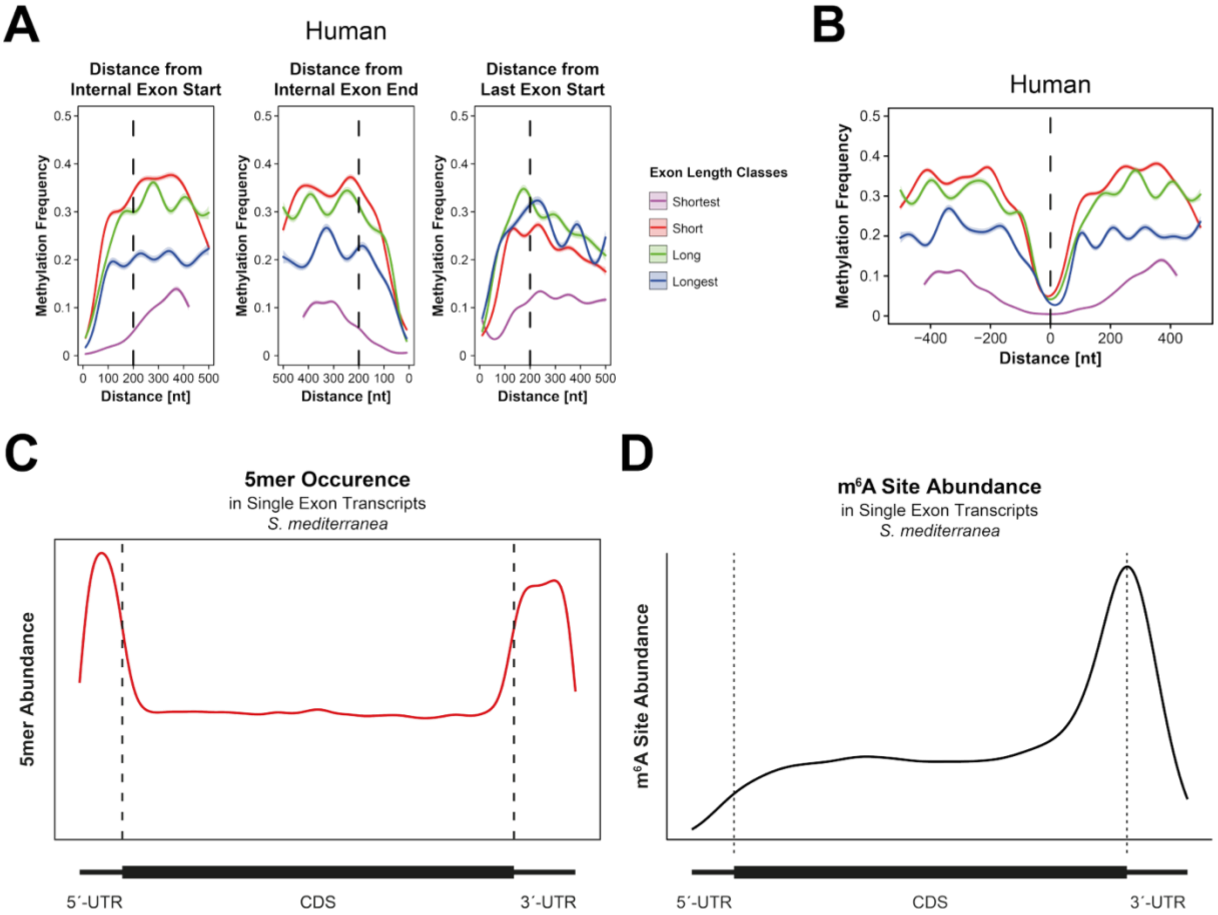


**Figure S2. Methylation frequency in human data recapitulates the EJC exclusion model. (A)** Generalized additive models (GAMs) of human methylation frequency towards exon boundaries reveal depletion of $m^6A$ in a ~200 nt window towards internal exon starts and ends, and last exon starts, recapitulating the $m^6A$ exclusion zone found in humans. GAMs were fitted for exon length classes defined by length quartiles. 95% confidence intervals of the model fit are depicted as shaded area; methylation frequency was tested in bins of 10 nt. **(B)** GAMs across human splice junctions recap the exclusion zone, which is centered on the splice junction and spans from ~ -200 nt to ~ +200 nt in humans. GAMs were fitted for exon length classes defined by length quartiles. 95% confidence intervals of the model fit are depicted as shaded area; methylation frequency was tested in bins of 10 nt. **(C-D)** Metagene plot of $m^6A$ sites in planarian single exon transcripts does not reveal a distinguished distribution of motif occurrences (C), however $m^6A$ site abundance in the CDS is elevated (D) compared to multi-exon transcripts (Fig. 1G, D)

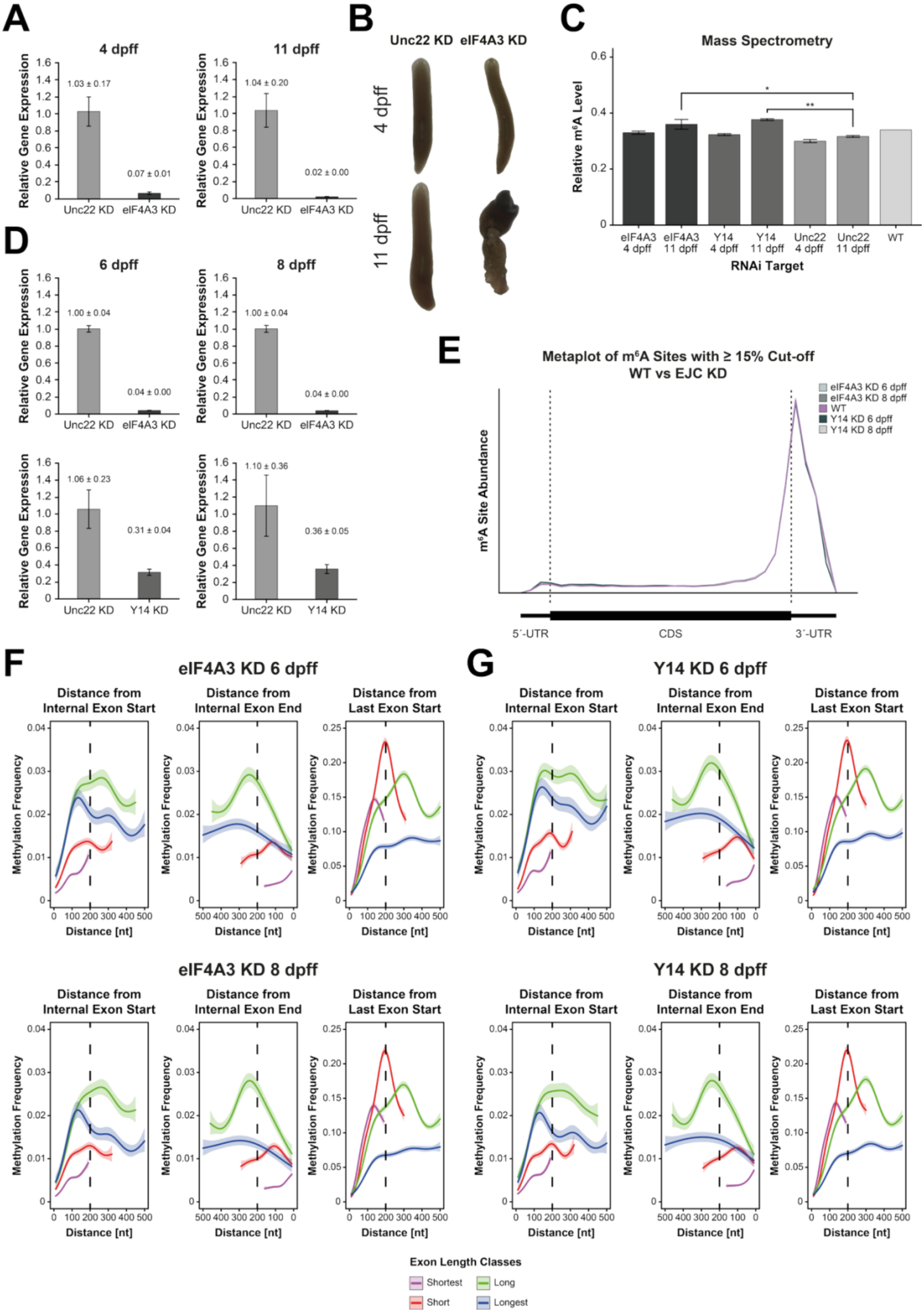
A
4 dpff
11 dpff
Relative Gene Expression
1.03 ± 0.17
0.07 ± 0.01
1.04 ± 0.20
0.02 ± 0.00
Unc22 KD
eIF4A3 KD
B
Unc22 KD
eIF4A3 KD
4 dpff
11 dpff
C
Mass Spectrometry
Relative m6A Level
eIF4A3 4 dpff
eIF4A3 11 dpff
Y14 4 dpff
Y14 11 dpff
Unc22 4 dpff
Unc22 11 dpff
WT
RNAi Target
D
6 dpff
8 dpff
1.00 ± 0.04
0.04 ± 0.00
1.06 ± 0.23
0.31 ± 0.04
1.10 ± 0.36
0.36 ± 0.05
Y14 KD
E
Metaplot of m6A Sites with ≥ 15% Cut-off
WT vs EJC KD
m6A Site Abundance
eIF4A3 KD 6 dpff
eIF4A3 KD 8 dpff
WT
Y14 KD 6 dpff
Y14 KD 8 dpff
5´-UTR
CDS
3´-UTR
F
eIF4A3 KD 6 dpff
eIF4A3 KD 8 dpff
G
Y14 KD 6 dpff
Y14 KD 8 dpff
Distance from Internal Exon Start
Distance from Internal Exon End
Distance from Last Exon Start
Methylation Frequency
Distance [nt]
Exon Length Classes
Shortest
Short
Long
Longest

**Figure S3. Validation of EJC component knockdowns and impact on exclusion zone. (A)** Validation of eIF4A3 knockdown in pre-experiments. Relative eIF4A3 mRNA levels were quantified by qPCR at 4 and 11 days post first feeding (dpff) following RNAi treatment. Efficient and sustained knockdown of eIF4A3 was observed compared to the Unc22 control. Y14 knockdown is not shown due to technical issues in the initial qPCR setup. Bars show mean ± SE. **(B)** Phenotypic consequences of eIF4A3 knockdown during planarian homeostasis. Representative images of animals subjected to RNAi against *unc22* or *eIF4A3* at 4 and 11 dpff. While control animals maintained normal morphology, *eIF4A3* knockdown resulted in pronounced homeostatic defects over time, supporting effective knockdown. Y14 knockdown animals are not shown. **(C)** Global $m^6A$ levels were quantified by mass spectrometry following RNAi-mediated depletion of the EJC components eIF4A3 and Y14 at the indicated time points. Relative $m^6A$ levels remained comparable to wild-type and Unc22 controls, indicating that EJC depletion does not lead to elevated global $m^6A$ levels. Bars show mean ± SE. Significance was tested using ANOVA and Tukey HSD test; p-values were corrected for multiple testing by Studentized range distribution; adjusted p-value: ** ≤ 0.01; * ≤ 0.05. **(D)** qPCR validation of *eIF4A3* and *y14* knockdown in samples used for nanopore sequencing. Relative eIF4A3 and Y14 mRNA levels were quantified at 6 and 8 dpff following RNAi treatment. Robust knockdown of both targets was achieved in these samples, which were subsequently used for nanopore direct RNA sequencing. Bars show mean ± SE. **(E)** Metagene analysis does not identify differential $m^6A$ site abundance profiles between wildtype and EJC depleted samples. **(F-G)** GAMs of methylation frequencies in *eIF4A3* (RNAi) and *y14* (RNAi) knockdowns towards exon boundaries at 6 dpff and 8 dpff did not reveal considerable differences in methylation frequencies compared to wildtype. GAMs were fitted for exon length classes defined by length quartiles in WT. 95% confidence intervals of the model fit are depicted as shaded area; methylation frequency was tested in bins of 10 nt.

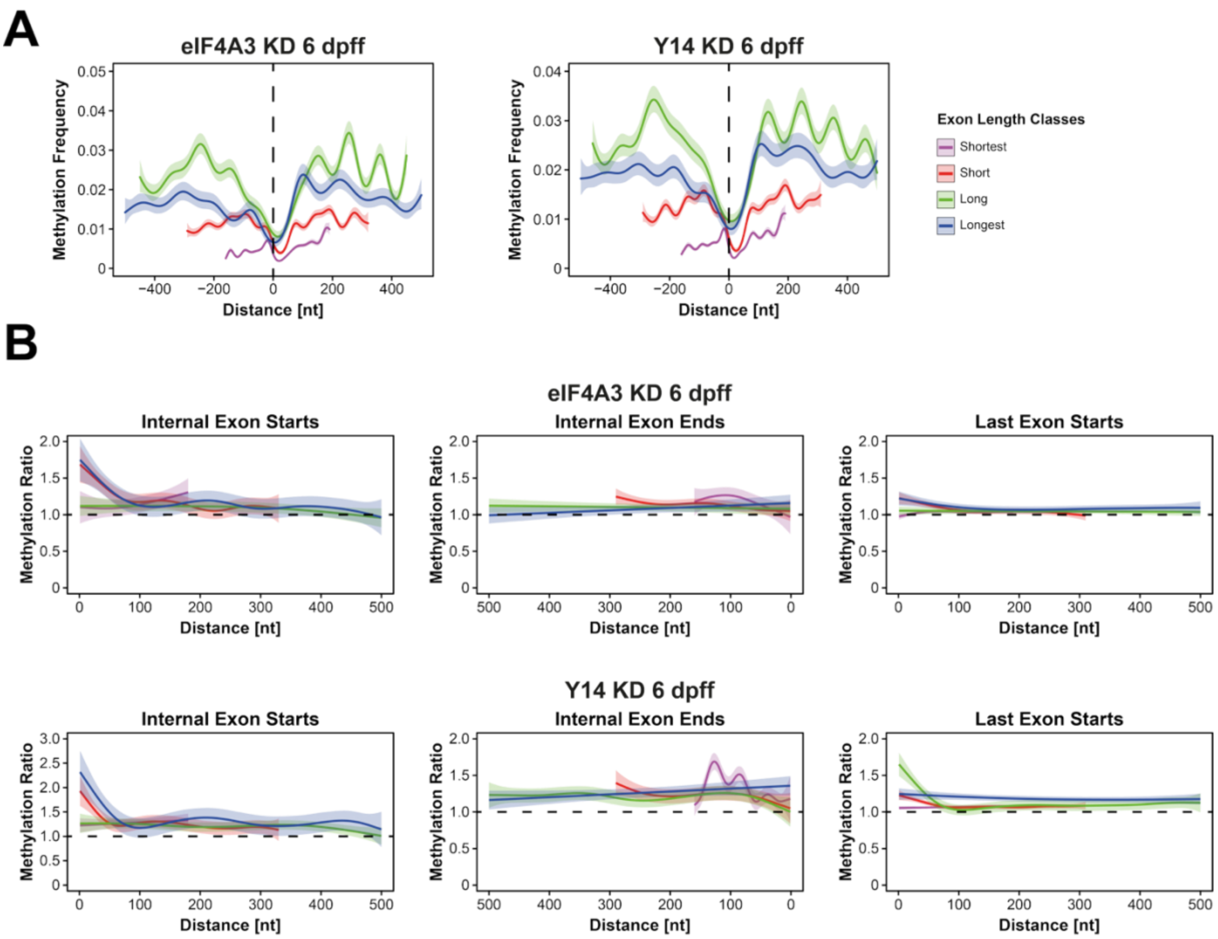


**Figure S4. Effects of EJC depletion are detectable at 6 dpff. (A)** GAMs of methylation frequency across splice junctions in *eIF4A3* (RNAi) and *y14* (RNAi) animals at 6 dpff reveal no considerable changes in $m^6A$ deposition compared to wild type (Fig. 2C). GAMs were fitted for exon length classes defined by length quartiles in WT. 95% confidence intervals of the model fit are depicted as shaded area; methylation frequency was tested in bins of 10 nt. **(B)** Generalized additive models (GAMs) of methylation frequency ratios between wild type and *eIF4A3* (RNAi) or *y14* (RNAi), respectively, at 6 dpff revealed an localized increase in methylation frequency at exon starts. GAMs were fitted for exon length classes defined by length quartiles in WT. 95% confidence intervals of the model fit are depicted as shaded area; methylation frequency was tested in bins of 10 nt.

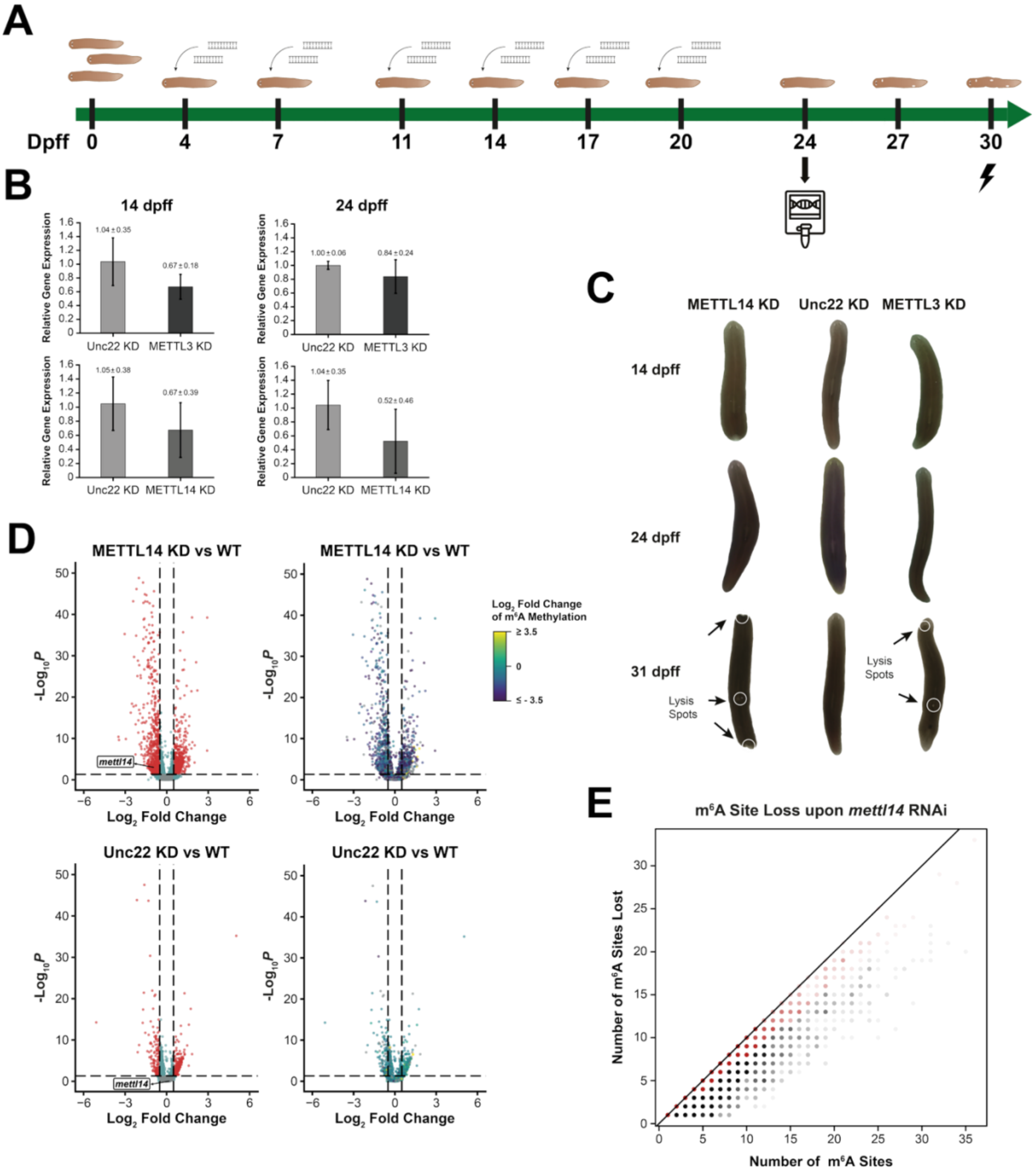
A
Dpff
0
4
7
11
14
17
20
24
27
30
B
14 dpff
24 dpff
Relative Gene Expression
Unc22 KD
METTL3 KD
METTL14 KD
1.04 ± 0.35
0.67 ± 0.18
1.00 ± 0.06
0.84 ± 0.24
1.05 ± 0.38
0.67 ± 0.39
1.04 ± 0.35
0.52 ± 0.46
C
METTL14 KD
Unc22 KD
METTL3 KD
14 dpff
24 dpff
31 dpff
Lysis Spots
Lysis Spots
D
METTL14 KD vs WT
METTL14 KD vs WT
Unc22 KD vs WT
Unc22 KD vs WT
-Log10P
Log2 Fold Change
mettl14
Log2 Fold Change of m6A Methylation
≥ 3.5
0
≤ - 3.5
E
m6A Site Loss upon mettl14 RNAi
Number of m6A Sites Lost
Number of m6A Sites

**Figure S5. Knockdown of MTC components via RNAi. (A)** Experimental timeline for METTL14 knockdown followed by nanopore dRNA sequencing. Animals were fed dsRNA at the indicated time points (days post first feeding, dpff) leading to phenotypical changes starting from 27 dpff. For nanopore dRNA sequencing, RNA of control and knockdown animals was extracted at 24 dpff. **(B)** Initial qPCR validation of METTL3 and METTL14 knockdown efficiency. Relative METTL3 and METTL14 mRNA levels were analyzed at 14 and 24 dpff following RNAi treatment. Both knockdowns resulted in reduced target gene expression compared to the Unc22 control, with METTL14 showing a stronger reduction at 24 dpff. Bars show mean ± SE. **(C)** MTC knockdown causes progressive defects during planarian homeostasis. Representative images of animals subjected to RNAi against METTL14, METTL3, or Unc22 at 14, 24, and 31 dpff. While control animals maintained normal morphology, METTL3 and METTL14 depletion led to progressive tissue defects and the appearance of lysis spots (indicated by arrows and white circles) at later stages. **(D)** Volcano plots of gene abundance changes between *mettl14* (RNAi) and wildtype, as well as *unc22* (RNAi) and wildtype conditions. Genes with a significant ($p \leq 0.05$) $log_2$-fold change $\leq -0.5$ or $\geq 0.5$ were considered as significantly deregulated (Left: red dots). On the right $m^6A$ $log_2$-fold change between respective conditions is displayed by dot color. **(E)** Scatterplot of the number of total vs lost $m^6A$ sites, with $m^6A$ sites being defined by a methylation rate ≥15% and loss of that site being defined as a reduction to 0%. We considered all transcripts that lost at least 80% of their $m^6A$ sites (red dots) for analysis with the Bayesian Mixture model.

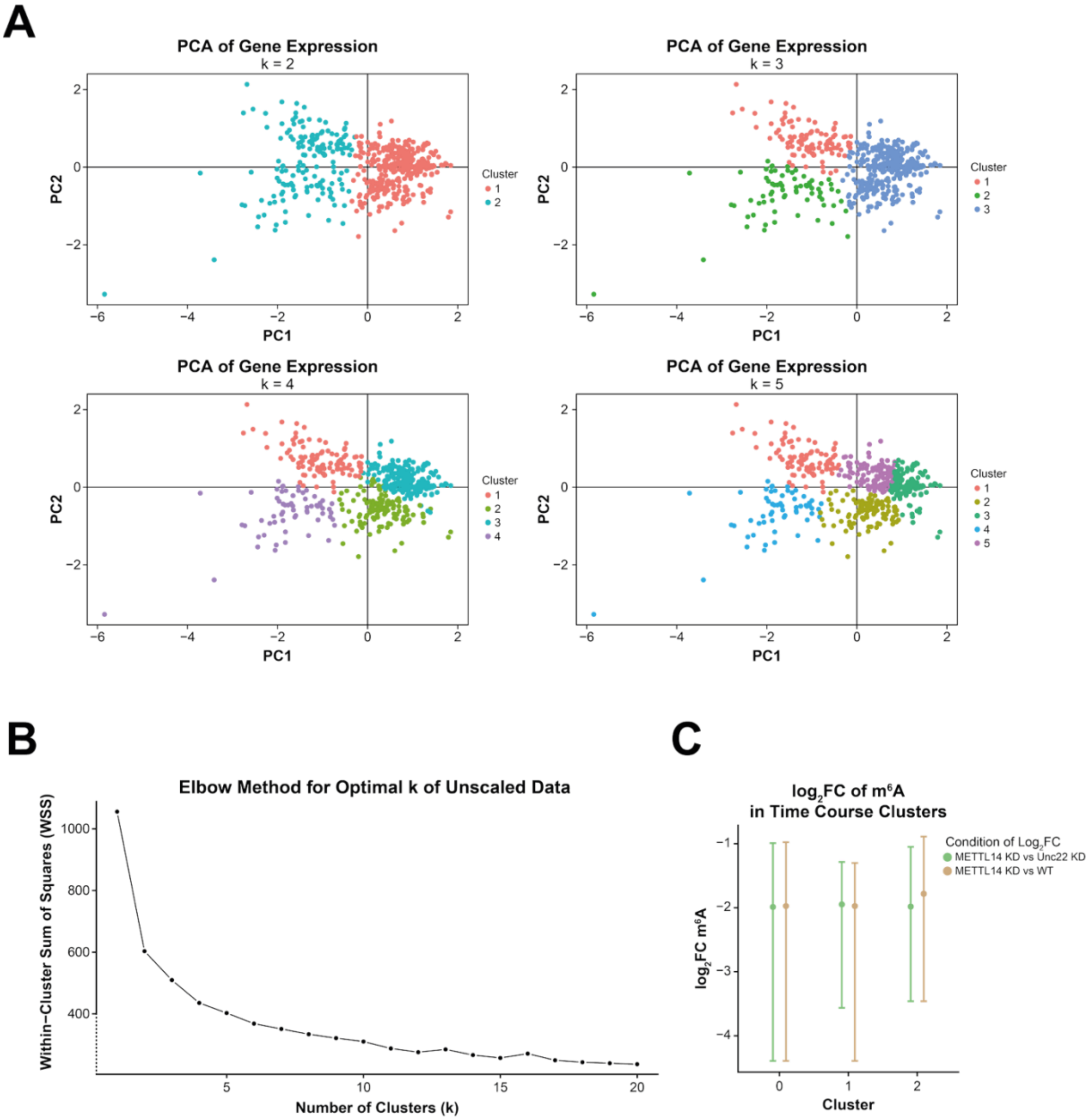


**Figure S6. Determination of cluster numbers in time series analysis. (A)** Principal component analysis plots for different cluster numbers displayed well defined clusters for k = 2, 3, and 4. **(B)** Within-cluster sum -of-squares (elbow-method) determined k = 2 as the optimal number of clusters for the time series analysis **(C)** Genes in time series clusters do not differ in loss of $m^6A$ sites compared to genes not clustered in wildtype conditions. Dots represent the median; whiskers mark the first and third quartile. Significance was tested using Kruskal-Wallis test and p-values were corrected for multiple testing with the Bonferroni method.
